# Preexisting memory CD4 T cells in naïve individuals confer robust immunity upon hepatitis B vaccination

**DOI:** 10.1101/2020.08.22.262568

**Authors:** George Elias, Pieter Meysman, Esther Bartholomeus, Nicolas De Neuter, Nina Keersmaekers, Arvid Suls, Hilde Jansens, Aisha Souquette, Hans De Reu, Evelien Smits, Eva Lion, Paul G. Thomas, Geert Mortier, Pierre Van Damme, Philippe Beutels, Kris Laukens, Viggo Van Tendeloo, Benson Ogunjimi

## Abstract

Antigen recognition through the T cell receptor (TCR) αβ heterodimer is one of the primary determinants of the adaptive immune response. Vaccines activate naïve T cells with high specificity to expand and differentiate into memory T cells. However, antigen-specific memory CD4 T cells exist in unexposed antigen-naïve hosts. In this study, we use high-throughput sequencing of memory CD4 TCRβ repertoire and machine learning to show that individuals with preexisting vaccine-reactive memory CD4 T cell clonotypes elicited earlier and higher antibody titers and mounted a more robust CD4 T cell response to hepatitis B vaccine. In addition, integration of TCRβ sequence patterns into a hepatitis B vaccine specific model can predict which individuals will have an early and more vigorous vaccine-elicited immunity. Thus, the presence of preexisting memory T clonotypes has a significant impact on immunity and can be used to predict immune responses to vaccination.

## Introduction

Antigen recognition through the T cell receptor (TCR) is one of the key determinants of the adaptive immune response (Rudolph et al., 2006). Antigen presentation via major histocompatibility complex (MHC) (encoded by HLA genes), together with the right costimulatory and cytokine signals, are responsible for T cell activation (Curtsinger and Mescher, 2010; Esensten et al., 2016). In this system, every T cell receptor (TCR) αβ heterodimer imparts specificity for a peptide-MHC (pMHC) complex. A highly diverse TCR repertoire ensures that an effective T cell response can be mounted against pathogen-derived peptides (Turner et al., 2009). High TCRαβ diversity is generated through V(D)J recombination at the complementary-determining region 3 (CDR3) of TCRα and TCRβ chains, accompanied with junctional deletions and insertions of nucleotides, further adding to the diversity (Krangel, 2009).

Vaccines activate naïve T cells with high specificity to vaccine-derived peptides and induce their expansion and differentiation into effective and multifunctional T cells. This is followed by a contraction phase from which surviving cells constitute a long-lived memory T cell pool that allows for a quick and robust T cell response upon a second exposure to the pathogen (Farber et al., 2014). However, recent work has shown that a prior pathogen encounter is not a prerequisite for the formation of memory T cells and that CD4 T cells with a memory phenotype can be found in antigen-naïve individuals (Su et al., 2013). The existence of memory-like CD4 T cells in naïve individuals (Sewell, 2012) can be explained by molecular mimicry, as the encounter with environmentally-derived peptides activates cross-reactive T cells due to the highly degenerate nature of the CD4 T cell recognition of peptide-MHC complex (Wilson et al., 2004). Indeed, work that attempted to replicate the history of human pathogen exposure in mice has shown that sequential infections altered the immunological profile and remodeled the immune response to vaccination (Reese et al., 2016). The existence of memory CD4 T cells specific to vaccine-derived peptides in unexposed individuals might confer an advantage in vaccine-induced immunity. In the present study we used high-throughput sequencing to profile the memory CD4 TCRβ repertoire of healthy adults before and after administration of a hepatitis B vaccine to investigate the impact of preexisting memory CD4 T cells on the immune response to the vaccine. Based on anti-hepatitis B surface (anti-HBs) antibody titers over 365 days, vaccinees were grouped into early, late and non-converters. Our data reveals that individuals with preexisting vaccine-specific CD4 T cell clonotypes in the memory CD4 compartment had earlier emergence of antibodies and mounted a more vigorous CD4 T cell response to the vaccine. Moreover, we identify a set of vaccine-specific TCRβ sequence patterns which can be used to predict which individuals will have an early and more vigorous response to hepatitis B vaccine.

## Results

### Vaccinee cohort can be classified into three groups

Out of 34 vaccinees, 21 vaccinees seroconverted (an anti-HBs titer above 10 IU/ml was considered protective (Keating and Noble, 2003)) at day 60 and were classified as early-converters; 9 vaccinees seroconverted at day 180 or day 365 and were classified as late-converters; remaining 4 vaccinees had an anti-HBs antibody titer lower than 10 IU/ml at all time points following vaccination and were classified as non-converters (**Fig. 1** and **Fig. S2a**).

**Figure 1.**
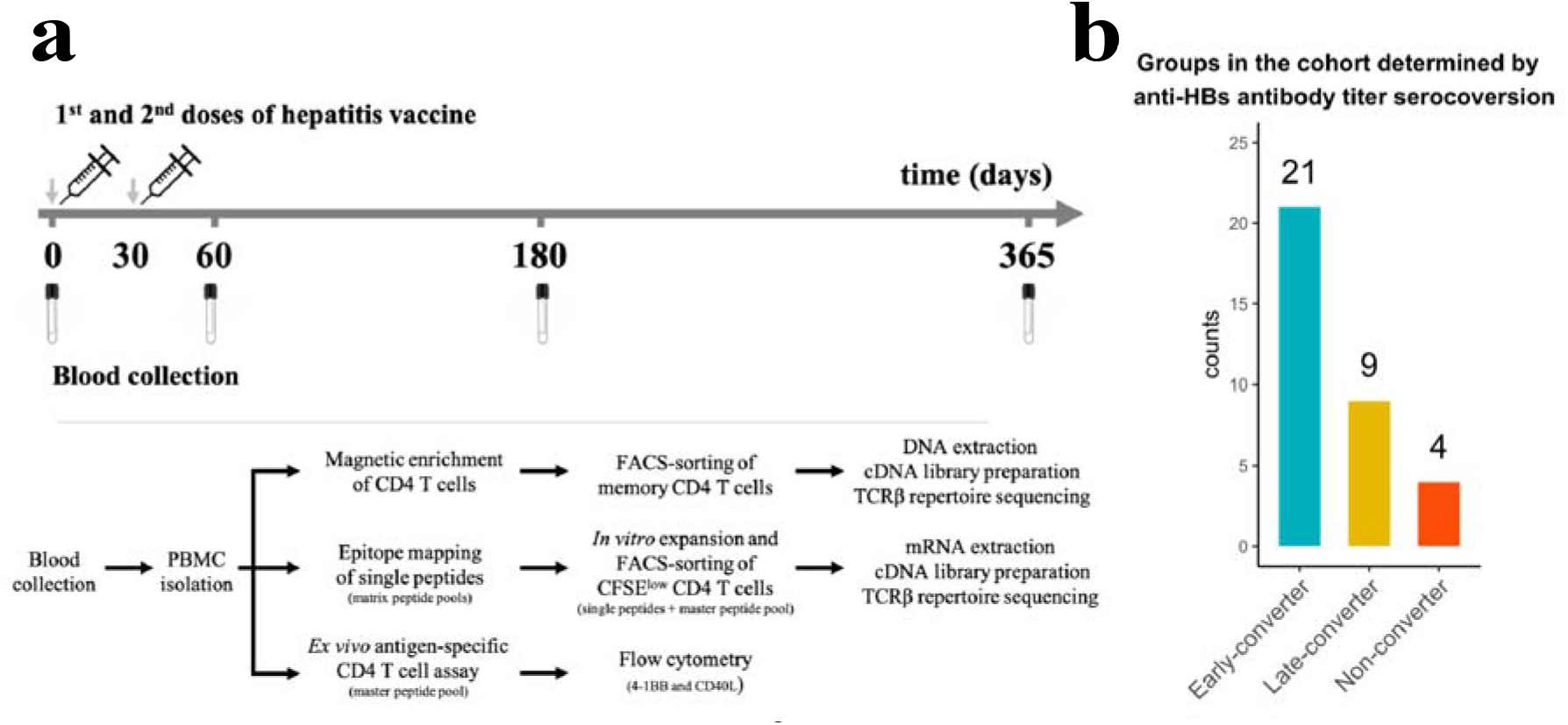
Hepatitis B vaccination (Engerix-B^®^) study design. **a** Hepatitis B (Engerix-B^®^) vaccination and experimental design. (Top) Timeline of vaccination and blood collection. (Bottom) Memory CD4 T cells were magnetically enriched and FACS-sorted from two time points (day 0 and day 60) for TCRβ vaccine from PBMCs repertoire sequencing. Matrix peptide pools were used to map CD4 T cell epitopes of the lected at day 60 and to select single peptides. After 7 days of *in vitro* expansion, single peptide-specific and master peptide pool-specific CFSE^low^ CD4 T cells from PBMCs collected at day 60 were FACS-sorted in two technical replicates for TCRβ repertoire sequencing. PBMCs collected at days 0, 60, 180, and 365 were stimulated with the master peptide HBsAg) and assessed for converse expression of 4-1BB and CD40L by flow cytometry. **b** Vaccinee cohort can be classified into three groups as determined by anti-Hepatitis B surface (anti-HBs) titer over four times points. Early-converters seroconverted at day 60, late-converters seroconverted at day 180 or day 365 and non–converters did not have an anti-HBs titer higher than 10 IU/ml at any of the time points.

Members of *Herpesviridae* family might alter immune responses to vaccines (Furman et al., 2015). We found no significant differences in CMV, EBV or HSV seropositivity between the three groups in our cohort (**Fig. S2b**). Early-converters were slightly younger than late-converters and non-converters were notably younger than both early and late converters (**Fig. S2c**).

### Memory CD4 T cell repertoire in early-converters decreases in clonality following vaccination

A genomic DNA-based TCRβ sequence dataset of memory CD4 T cells isolated from peripheral blood was generated from a cohort of 33 healthy vaccinees (see *Methods* for details) right before vaccination (day 0) and 60 days after administration of the first dose of hepatitis B vaccine (30 days after administration of the second vaccine dose).

Between 4.54 × 10^4^ and 3.92 × 10^5^ productive TCRβ sequence reads were obtained for each vaccinee at each time point (**Fig. S3a**). Between 30,000 and 90,000 unique TCRβ sequences were sequenced for each vaccinee at each time point (**Fig. S3b**). As expected, considering the extremely diverse memory CD4 T cell repertoire (Klarenbeek et al., 2010), less than 20% of the TCRβ sequences is shared between the time points for each vaccinee (**Fig. S3c**).

The diversity of the memory CD4 T cell repertoire of each vaccinee at the two time points was explored. Even though the number of unique clones in the memory CD4 TCRβ repertoire remained stable in between the two time points, we detected a significant increase in the TCRβ repertoire Shannon’s entropy for early-converters (**Fig. 2a**) (P value = 0.042), but not late-converters, suggesting that the memory CD4 T cell repertoires of early-converters have become less clonal, despite the number of distinct TCRβ sequences not changing significantly.

**Figure 2.**
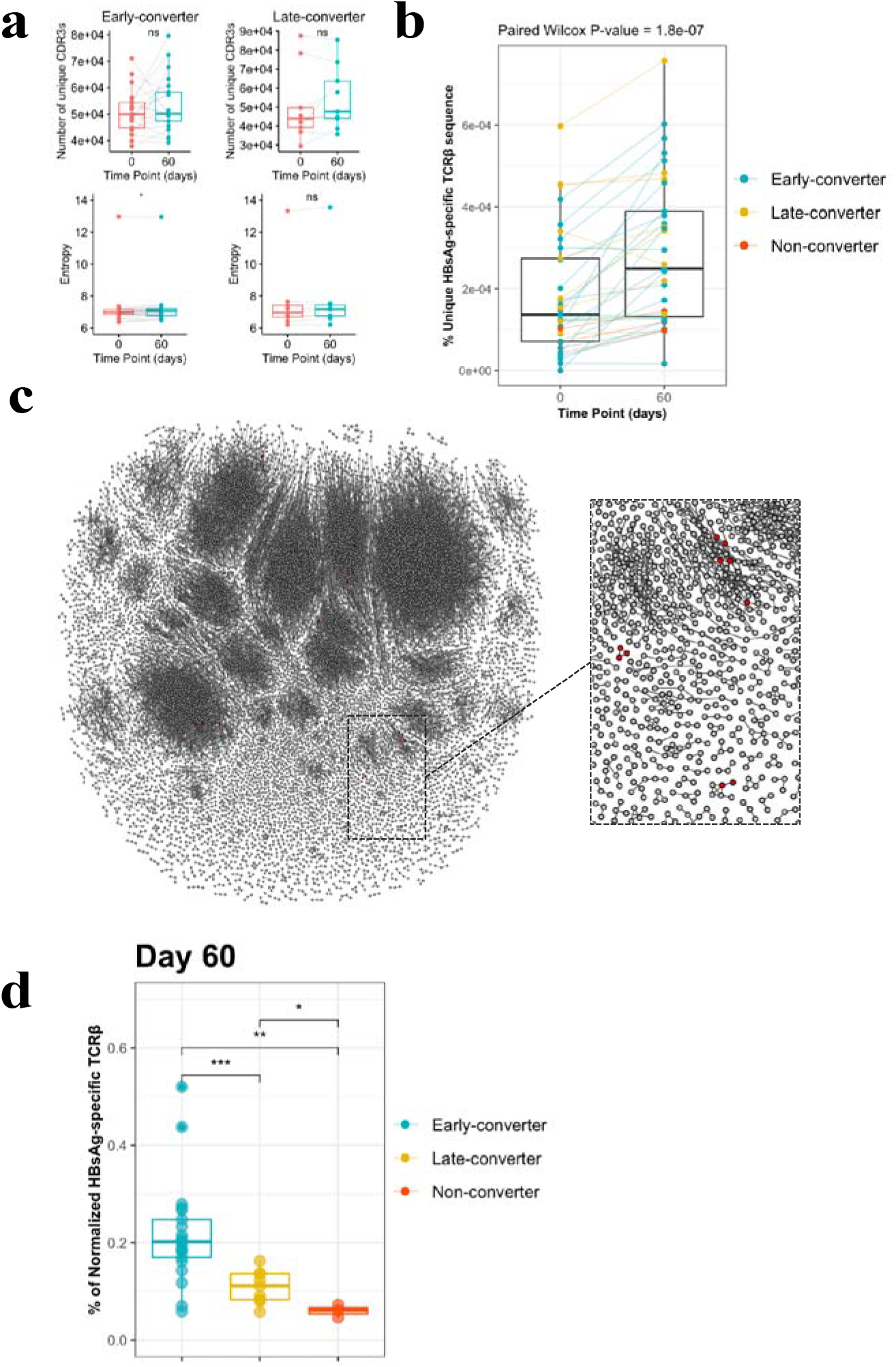
CD4 T cell memory TCRβ repertoire and vaccine-specific TCRβ clonotypes. a Comparison of the memory CD4 T repertoire diversity and entropy between day 0 and day 60. b Frequency of unique vaccine-specific TCRβ sequences out of total sequenced TCRβ sequences between two time points for all vaccinees colored by group. c Sequenced CD4^+^ TCR memory repertoire of vaccinee H35 at day 60. Each TCR clonotype is represented by a node. TCRs are connected by an edge if their Hamming distance is one. Only clusters with at least three TCRs are shown. TCR clonotypes in red are the vaccine-specific TCRβ sequences that were not present prior to vaccination. d Frequency of vaccine-specific TCRβ sequences within memory CD4 T cell repertoire normalized by number of HBsAg-specific TCRβ sequences found for each vaccinee at time point 60.

### Unique vaccine-specific TCRβ sequences are trackable within memory CD4 T cell repertoire and increase following vaccination

Peripheral blood mononuclear cells from day 60 were labeled with carboxyfluorescein succinimidyl ester (CFSE) and stimulated with a pool of peptides spanning hepatitis B (HB) surface antigen (HBsAg). After day 7 of *in vitro* expansion, we sorted CFSE^low^ CD4 T cells (Becattini et al., 2015) and extracted mRNA for quantitative assessment of HBsAg-specific TCRβ clonotypes by sequencing (see *Methods* for details), allowing for the tracking of vaccine-specific TCRβ within memory CD4 T cell repertoire over the two times points, based on CDR3β amino acid sequence mapping.

We detected a significant increase in the frequency of unique HBsAg-specific TCRβ sequences at day 60 post-vaccination compared to pre-vaccination (mean increase = 96.5%, 95% CI = 56.7 - 170%) (**Fig. 2b** and **S3d**). Moreover, this increase was larger for early-converters (mean = 132.1%, 95% CI = 76.4 - 238.2%) than late-converters (mean = 22.1%, 95% CI = 5.9 – 50.1%).

For non-converters the mean was 81.6% (95% CI: [42.7% - 110.6%]). A Wilcoxon test shows that the difference between the increase for the early converters and late converters had a P value of 0.04909.

As HBsAg-specific TCRβ sequences were already detected in the memory CD4 T cell repertoire prior to vaccination, we sought to determine whether the vaccination results in an expansion of those sequences. Using the abundance of vaccine-specific TCRβ sequences within the memory CD4 T cell repertoire, the data does not support a vaccine-induced expansion of preexisting vaccine-specific TCRβ sequences (**Fig. S3e**). Thus, although we see a rise in the number of vaccine-specific TCRβ clonotypes from day 0 to day 60, this cannot be attributed to an expansion of preexisting TCRβ clonotypes but rather the recruitment of new TCRβ clonotypes (presumably from the naïve T cell compartment), as visualized for one vaccinee in **Fig. 2c**.

It makes sense to not only look at the difference in vaccine-specific TCRβ sequences between time points, but also explore whether there are differences in the proportion of HBsAg-specific clones in the memory repertoire between early-converters, late-converters and non-converters after vaccination. In this case, as we aim for a between-vaccinees comparison (in contrast to the within-vaccinees timepoint comparison), we normalize by the number of HBsAg-specific TCRβ found for each vaccinee. Thus, the values are different from those reported before. From this analysis, it can be concluded that there is a difference in HBsAg-specific TCRβ at day 60 between the three groups (**Fig. 2d**) (ANOVA P value = 0.00238). A Wilcoxon test between early-converters and other vaccinees shows a significant P value of 0.000473, indicating that early-converters have a higher relative frequency of vaccine-specific TCRβ sequences present in their memory CD4 T cell repertoire at day 60 compared to vaccinees from the two other groups in the cohort.

### HBsAg single peptide-specific TCRβ identification allows predictive modelling of early converters prior to vaccination

To quantify the T cell response at the level of individual peptides that make up the HBsAg, a matrix peptide pool covering 54 overlapping peptides of the HBsAg was used to extract peptide-specific T-cells using a CD40L/CD154 activation-induced marker (AIM) assay (see *Methods* for details, **Fig. S4**). The top 6 peptides for each individual were selected for TCR sequencing after a CFSE assay (**Supplementary Table 2**). In this manner, TCRβ sequences were identified for T-cells reactive against 44 single HBsAg peptides. These were not uniformly distributed across the HBsAg amino acid sequence, with the most prominent epitopes covering the regions 1-15, 129-144, 149-164, 161-176, 181-200, 213-228. For each of those regions, more than 10 individuals had a strong T-cell response and more than 150 unique TCRβ sequences could be identified **(Fig. 3a)**.

**Figure 3.**
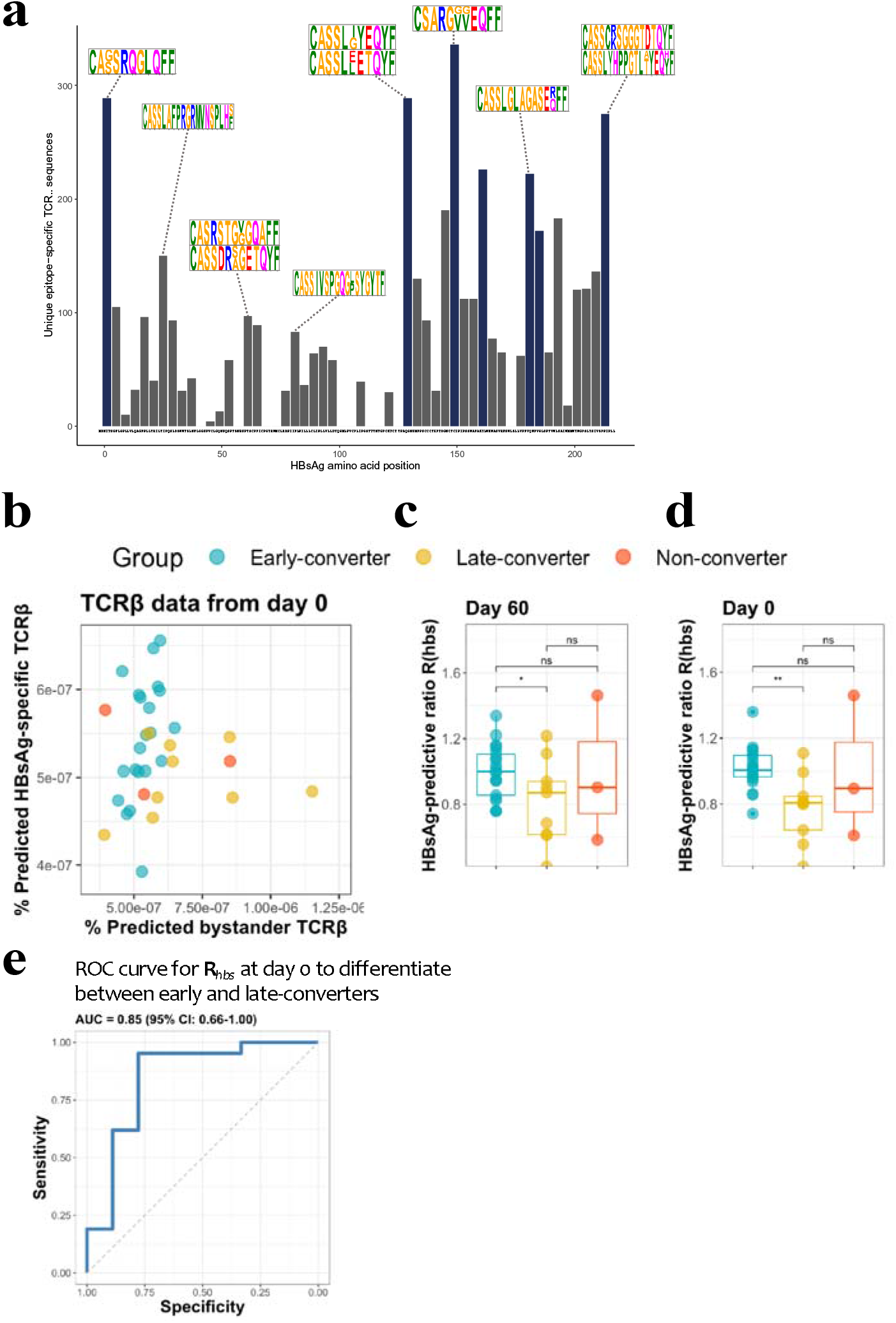
HBsAg peptide-specific TCRβ identification and predictive potential of R*_hbs_*. **a** Overview of the detected HBsAg epitope-specific TCRβ sequences. Each bar corresponds to unique TCRβ sequences found against a single 15mer HBsAg peptide, with 11 amino acid overlap to each subsequent peptide. Bars in blue denote those epitopes for which 10 or more volunteers had a strong T-cell reaction. Motif logos on top of bars denote a sampling of the most common TCRβ amino acid sequence motifs for those epitopes. **b** Scatter plot with the frequency of predicted HBsA epitope-specific and bystander TCRβ sequences. Predictions done as a leave-one-out cross-validation. Each circle represents a vaccinee with the color denoting the response group (blue: early-converter, yellow: late-converter, red: non-converter). **c** HBsAg-predictive ratio, **R***_hbs_*, when calculated on the memory CD4 TCR repertoires at day 60. **d** HBsAg-predictive ratio, **R***hbs*, when calculated on the memory CD4 TCRβ repertoires at day 0. **e** Receiver operating characteristic (ROC) curve using **R***_hbs_* to differentiate ^β^ tween early-converters and late-converters in a leave-one-out cross validation at day 0. Reported is the area under the curve (AUC) and its 95% confidence interval.

These peptide-specific TCRβ sequences can be utilized in a peptide-TCR interaction classifier to identify other TCRβ that are likely to react against same HBsAg epitopes, as it has been shown that similar TCRβ sequences tend to target the same epitopes (Meysman et al., 2018; De Neuter et al., 2018). These classifications were integrated into a model which outputs a **ratio R***_hbs_* for any TCRβ repertoire representing the amount of HBsAg peptide-specific clonotypes. **R***_hbs_* is based on the frequency of putative peptide-specific TCRβ divided by a normalization term for putative false positive predictions due to bystander activations in the training data set. This model applied to the memory repertoire at day 60 shows that early-converters tend to have a higher frequency of putative HBsAg peptide-specific TCRβ, while late-converters tend to have relatively more false positive hits (**Fig. 3b**). Thus, the defined ratio **R***_hbs_* shows significant difference between early the late-converters at day 60 (one-sided Wilcoxon-test P value= 0.0313, **Fig. 3c**). Furthermore, calculating **R***_hbs_* on the memory repertoires prior to vaccination (day 0) shows a similar difference (one-sided Wilcoxon-test P value= 0.0010, **Fig. 3d**). In this manner, **R***_hbs_* has predictive potential and can be used as a classifier to distinguish early from late-converters prior to vaccination (**Fig. 3e**), with an AUC of 0.825 (95% CI: 0.657 – 0.994) in a leave-one-out cross validation setting. To account for the age variable, a model in which age-matched vaccinees were included from early and late-converters returned a similar ROC curve (**Fig. S3f**).

While **R***_hbs_* is able to differentiate between early-and late-converters, it seems to be worse at distinguishing non-converters. This is mainly due to a single non-converter vaccinee (H21) with a high **R***_hbs_*, signifying a high number of putative HBsAg peptide-specific TCRβ in their memory repertoire.

### Vaccine-specific conventional and regulatory memory CD4 T cells induced in early-converters

After showing evidence for the existence of vaccine-specific TCRβ sequences pre-vaccination and that individuals with a higher number of HBsAg peptide-specific clonotypes had earlier seroconversion, we attempted to link this observation to differences in vaccine-specific CD4 T cells responses using CD4 T cell assays. As T_REG_ cells might suppress vaccine-induced immune responses (Brezar et al., 2016), we used activation markers CD40L (CD154) and 4-1BB (CD137) to help delineate the conventional (T_CON_) and regulatory (T_REG_) phenotypes of activated CD4 T cells. In this scheme, after 6 hours of antigen stimulation, CD40L^+^4-1BB^−^ can be used as a signature for antigen-specific CD4 T_CON_ cells, as opposed to CD40L^−^4-1BB^+^ signature for antigen-specific CD4 T_REG_ cells (Elias et al., 2020; Schoenbrunn et al., 2012).

Additionally, we added CD25 and CD127 to better identify T_REG_ cells (Liu et al., 2006; Seddiki et al., 2006) and CXCR5 to further distinguish circulatory T follicular helper cells (cT_FH_) and circulatory T follicular regulatory cells (cT_FR_) (Bentebibel et al., 2011; Fonseca et al., 2017). Using the converse expression of CD40L and 4-1BB, CD40L^+^4-1BB^−^ and CD40L^−^4-1BB^+^ CD4 T cells had a T_CON_ and T_REG_ phenotype, respectively, as shown by the expression of CD25 and CD127 (**Fig. S5a** and **b**), and validate their use for the distinction of activated T_CON_ and T_REG_ cells as has been reported before (Schoenbrunn et al., 2012).

We detected a significant increase in the frequency of CD40L^+^4-1BB^−^ and CD40L^−^4-1BB^+^ memory CD4 T cells at day 60 in our cohort (**Fig. 4a**) that correlated positively with the increase in antibody titer between day 0 and day 365 (**Fig. 4b** and **Fig. S6**). Upon a closer look, the induction of both signatures of vaccine-specific memory CD4 T cells was only true for early-converters (**Fig. 4c**, see **Fig. S7a** for non-converters and **Fig. S7b** for vaccine-specific CD4 T cells) while late-converters did not show a detectable memory CD4 T cell response. Although a subset of both early and late-converters had detectable memory CD4 T cell responses prior to vaccination, we observed no significant differences in the frequencies of CD40L^+^4-1BB^−^ and CD40L^−^4-1BB^+^ memory CD4 T cells between the two groups at day **0** (**Fig. 4d**).

**Figure 4.**
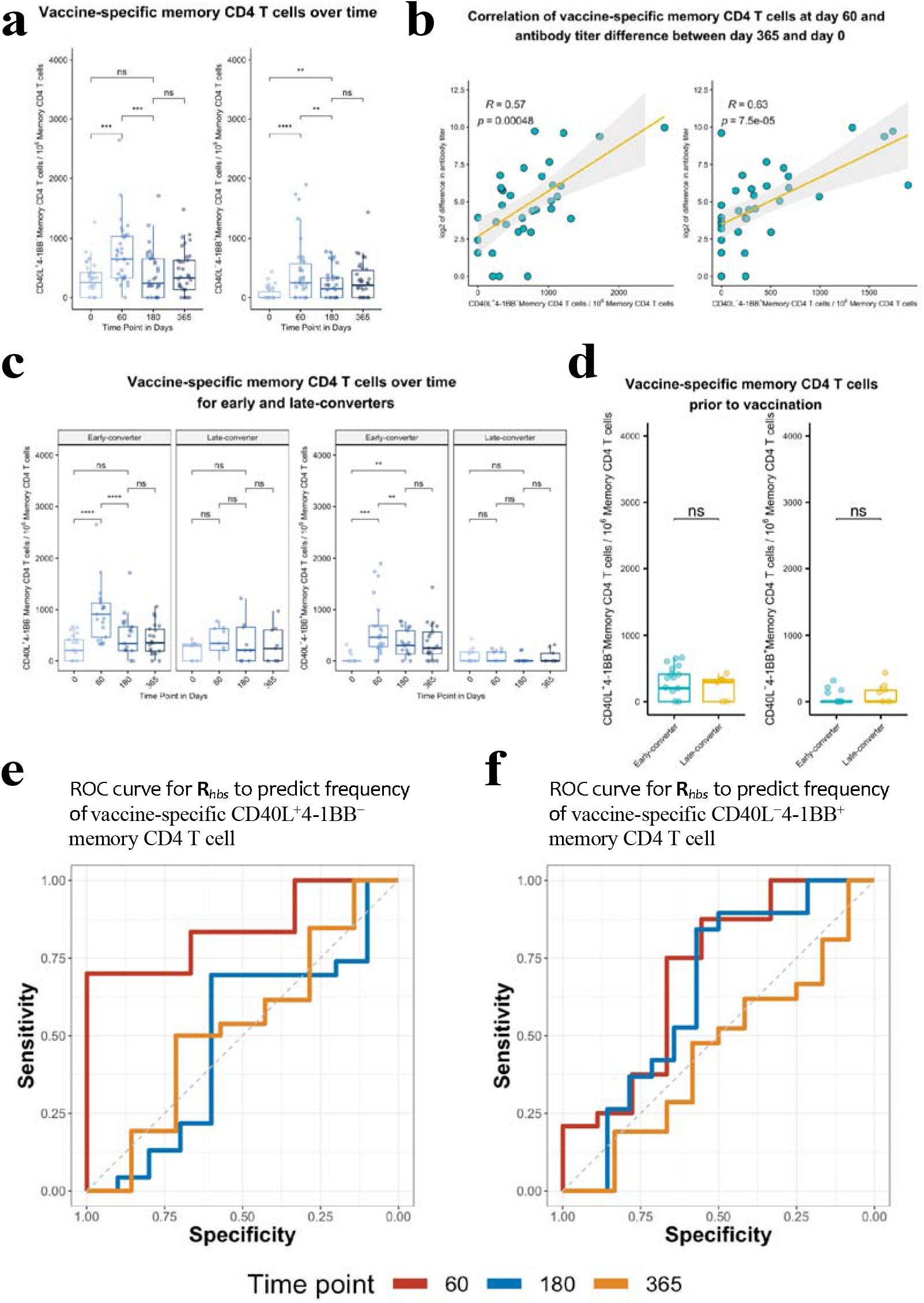
Hepatitis B vaccine induces a vaccine-specific CD40L^+^4-1BB^−^ and CD40L^−^4-1BB^+^ memory CD4 T cell response in early-converter vaccinees. PBMCs from vaccinees were stimulated with 2 μg/ml of the master peptide pool (HBsAg) and assessed for converse expression of 4-1BB and CD40L by flow cytometry on days 0, 60, 180, and 365. Shown is number of of vaccine-specific memory CD4 T cells out of 10^6^ memory CD4 T cells after subtraction of responses in negative control. **a** Aggregate analysis from vaccinees (including early, late and non-converters) showing a peak of vaccine-specific CD40L^+^4-1BB^−^ and CD40L^−^4-1BB^+^ memory CD4 T cell at day 60 (day 60 after 1^st^ dose of the vaccine and day 30 after 2^nd^ dose), declining thereafter. Shown are numbers of vaccine-specific memory CD4 T cells out of 10^6^ memory CD4 T cells. **b** Correlation between the difference in antibody titer between day 365 and day 0 and vaccine-specific CD40L^+^4-1BB^−^ and CD40L^−^4-1BB^+^ memory CD4 T cell at day 60. **c** Aggregate analysis from early and late-converter vaccinees showing a significant induction of vaccine-specific CD40L^+^4-1BB^−^ and CD40L^−^4-1BB^+^ memory CD4 T cell in early-converters and lack thereof in late-converters. **d** Aggregate analysis from early and late-converter vaccinees showing no significant differences in vaccine-specific CD40L^+^4-1BB^−^ and CD40L^−^4-1BB^+^ memory CD4 T cell at day 0. **e** Receiver operating characteristic (ROC) curves for R*_hbs_* from day 0 data in a leave-one-out cross-validation compared to the frequency of vaccine-specific CD40L^+^4-1BB^−^ memory CD4 T cell out of 10^6^ memory CD4 T cells for each vaccinee at time points 60 (AUC = 0.84), 180 (AUC = 0.56) and 365 (AUC = 0.57). **f** Receiver operating characteristic (ROC) curves for R*_hbs_* from day 0 data in a leave-one-out cross-validation compared to the frequency of vaccine-specific CD40L^−^4-1BB^+^ memory CD4 T cell out of 10^6^ memory CD4 T cells for each vaccinee at time points 60 (AUC = 0.62), 180 (AUC = 0.56) and 365 (AUC = 0.52). Wilcoxon signed-rank with unpaired and paired analysis as necessary; statistical significance was indicated with ns P > 0.05, * P ≤ 0.05, ** P ≤ 0.01, *** P ≤ 0.001, **** P ≤ 0.0001 *rs*, Spearman correlation coefficient, −1 ≤ *r*s ≤ 1; *rs* and *p* value by Spearman’s correlation test

Collectively, flow cytometry data reveal that the expression of CD40L and 4-1BB in our ex vivo assay is consistent with our serological data and reflects the lack of seroconversion at day 60 in late-converters. However, it does not support the existence of more vaccine-specific memory CD4 T cells in early-converters prior to vaccination.

### Predictive capacity of TCRβ repertoire holds true for CD4 T_CON_ immune response

Response groups used thus far were established based on the dynamics of anti-HBs titers following vaccination. However, response groups can be defined based on the data of antigen-specificity from the ex vivo CD4 T cell assay. Redoing the analysis with **R***_hbs_* to predict the frequency of CD40L^+^4-1BB^−^ and CD40L^−^4-1BB^+^ memory CD4 T cells at three different times points (days 60, 180 and 365) shows that the **R***_hbs_* is a good classifier in a leave-one-out cross validation for HBsAg-specific memory CD4 T cells with a T_CON_ signature identified at day 60 post-vaccination (**Fig. 4e** and **f**) and that is to a large extent lacking for delayed time points.

### An expanded subset of 4-1BB^+^CD45RA^−^ T_REG_ cells is a prominent feature of late-converters

In order to detect any distinct signatures of early and late-converters, we analyzed pre-vaccination flow cytometry data to examine major CD4 T cell subsets; T_H_, T_REG_, cT_FH_ and cT_FR_ cells. Using manual gating in which regulatory T cells (T_REG_) were defined as viable CD3^+^CD4^+^CD8^−^CD25^+^CD127^−^CXCR5^−^ and were further divided into CD45RA^+^ and CD45RA^−^ T_REG_ cells, we identified a significantly higher frequency of 4-1BB^+^ CD45RA^−^ T_REG_ cells in late-converters compared to early-converters **(Fig. 5a and Fig. S8)**.

**Figure 5.**
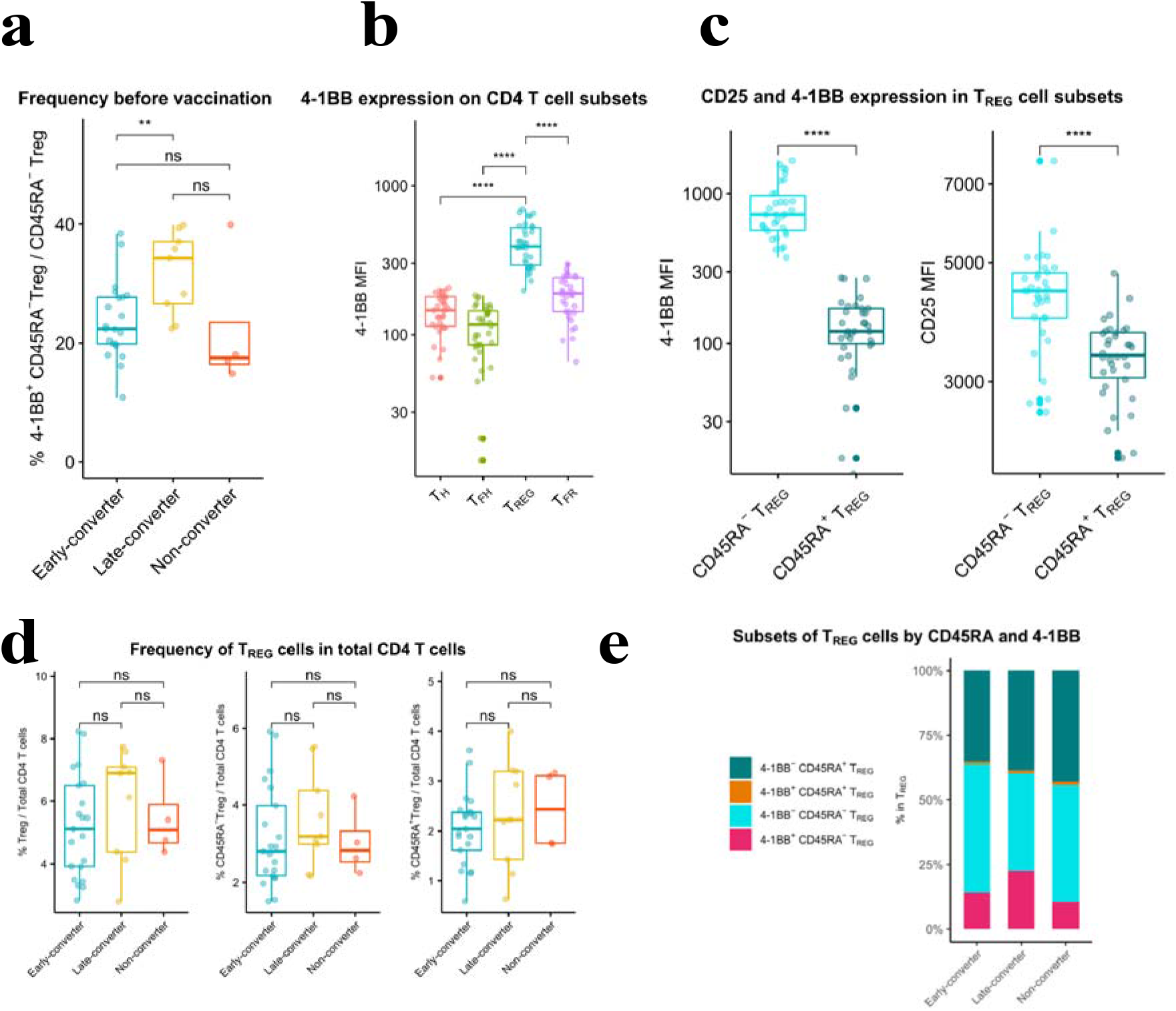
An expanded 4-1BB^+^CD45RA^−^ T_REG_ cells within T_REG_ compartment is a prominent feature in late-converters prior to vaccination. PBMCs from vaccinees at day 0 (prior to vaccination) were phenotyped for expression of markers of T_REG_. **a** Aggregate analysis of 4-1BB^+^CD45RA^−^ T_REG_ within CD45RA^−^ T_REG_ CD4 T cells in early and late and non-converter vaccinees before vaccination. **b** Aggregate analysis of the median fluorescence intensity of 4-1BB in T_H_, cT_FH_, T_REG_ and cT_FR_ cells before vaccination. **c** Aggregate analysis of the median fluorescence intensity of 4-1BB (left panel) and CD25 (right panel) in CD45RA^−^ T_REG_ and CD45RA^+^ T_REG_ cells before vaccination. **d** Frequency of T_REG_, CD45RA^−^ T_REG_ and CD45RA^+^ T_REG_ cells within total CD4 T cells in early, late and non-converter vaccinees before vaccination. **e** Composition of T_REG_ compartment as determined by expression of 4-1BB and CD45RA in early, late and non-converter vaccinees before vaccination. Wilcoxon signed-rank with unpaired and paired analysis as necessary; statistical significance was indicated with ns P > 0.05, * P ≤ 0.05, ** P ≤ 0.01, *** P ≤ 0.001, **** P ≤ 0.0001

T_REG_ cells showed higher 4-1BB expression compared to T_H_, cT_FH_ and cT_FR_ cells **(Fig. 5b)** and within T_REG_ subset, CD45RA^−^ T_REG_ cells showed significantly higher expression of 4-1BB, accompanied with a higher expression of CD25, compared to CD45RA^+^ T_REG_ cells **(Fig. 5c)**. In this scheme, CD45RA^−^ T_REG_ can be divided into 4-1BB^+^CD25^high^ and 4-1BB^−^CD25^int^ subsets. It is worth noting here that no differences were detected in the frequency of CD45RA^−^ or CD45RA^+^ T_REG_ cells within CD4 T cell compartment between the two groups **(Fig. 5d)**, and that the composition of T_REG_ compartment that is distinct between the two groups **(Fig. 5e)**.

In summary, an expanded subset of 4-1BB^+^CD45RA^−^ T_REG_ cells pre-vaccination is a prominent feature of a delayed and modest immune response to hepatitis B vaccine in our cohort.

## Discussion

In this study, we used high-throughput TCRβ repertoire profiling and *ex vivo* T cell assays to characterize memory CD4 T cell repertoires before and after immunization with hepatitis B vaccine, an adjuvanted subunit vaccine, and tracked vaccine-specific TCRβ clonotypes over two time points. As antigen-naïve adults were found to have an unexpected abundance of memory-phenotype CD4 T cells specific to viral antigens (Su and Davis, 2013; Su et al., 2013), we sought to investigate the influence that preexisting memory CD4 T cells can have on vaccine-induced immunity.

Commercially available HBV vaccines produces a robust and long-lasting anti-HBs response, and protection is provided by induction of an anti-HBs (antibody against HBV surface antigen) titer higher than 10 mIU/mL after a complete immunization schedule of 3 doses (Meireles et al., 2015). However, 5-10% of healthy adult vaccinees fail to produce protective titers of anti-HBs and can be classified as non-responders (Meireles et al., 2015). In our cohort, 13 vaccinees did not seroconvert by day 60 (30 days following administration of the second vaccine dose), as determined by antibody titer. Out of this group, 9 vaccinees seroconverted by day 180 or day 365, referred to here as late-converters, and 4 vaccinees did not seroconvert, referred to here as non-converters.

A hallmark of adaptive immunity is a potential for memory immune responses to increase in both magnitude and quality upon repeated exposure to the antigen (Sallusto et al., 2010). Our systems immunology data supports the theory that preexisting memory CD4 T cell TCRβ sequences specific to HBsAg, the antigenic component of the current hepatitis B vaccine, predict which individuals will mount an early and more vigorous immune response to the vaccine as evidenced by a higher fold change in anti-HBs antibody titer and a more significant induction of antigen-specific CD4 T cells. It is postulated that preexisting memory CD4 T cell clonotypes are generated due to the highly degenerate nature of T cell recognition of antigen/MHC and are cross-reactive to environmental antigens (Sewell, 2012). For example, preexisting memory CD4 T cells are well-established in unexposed HIV-seronegative individuals, although at a significantly lower magnitude than HIV-exposed seronegative individuals (Campion et al., 2014; Ritchie et al., 2011), and were likely primed by exposure to environmental triggers or the human microbiome.

We and others have shown before that the TCRβ repertoire of CD4 T cells encodes the antigen exposure history of each individual and that antigen-specific TCRβ sequences could serve to automatically annotate the infection or exposure history (DeWitt et al., 2018; Emerson et al., 2017; de Neuter et al., 2018). In this study, we show that similar principles can be used to study vaccine responsiveness. Specifically, the recruitment of novel vaccine-specific T-cell clonotypes into memory compartment following vaccination can be tracked by examining the CD4 memory TCRβ repertoire over time. While we observed no increase in the frequency of the vaccine-specific memory T-cells, as the time point may have missed the peak of the clonal expansion of effector CD4 T cells as was reported before (Blom et al., 2013; Kohler et al., 2012; Pogorelyy et al., 2018), a significant rise in the number of unique vaccine-specific T-cell clonotypes was detected. This observation is consistent with earlier studies of T cell immune repertoire that showed that antigen-specific TCRβ sequences do not always overlap with those sequences that increase in frequency after infection or vaccination (DeWitt et al., 2015). More interestingly, individuals with the earlier and more robust response against the vaccine, had a telltale antigen-specific signature in their memory TCRβ repertoire prior to vaccination, despite the lack of HBsAg antibodies or prior vaccination history.

Detection of this vaccine-specific signature was possible due to the development of a novel predictive model that used epitope-specific TCRβ sequences from one set of individuals to make predictions about another. A correction factor was needed to account for the occurrence of bystander activated T cells within the original epitope-specific TCRβ sequences. Indeed, in those vaccinees without a positive antibody titer at day 60, putative vaccine-specific T cells might be induced duo to bystander activation. This was supported by predictions using the TCRex tool (Gielis et al., 2019), which matched these TCR sequences to common viral or other epitopes. It is of note that these TCRβ sequences are matched with CD8 T cell epitopes, while they originate from isolated CD4 T cells. This is likely due to the great similarity between the TCRβ sequences of CD4 and CD8 T cells as noted in prior research (Meysman et al., 2018).

However, our in vitro antigen-specificity data, using an assay that enables discrimination of T_CON_ and T_REG_ cells using the converse expression of the activation markers CD40L and 4-1BB (Frentsch et al., 2005; Schoenbrunn et al., 2012), failed to show a significant difference in preexisting antigen-specific CD4 T cells between early and late-converters prior to vaccine administration. It is plausible that the signal is below the detection limit of the assay and that more sensitive assays that require pre-enrichment of CD40L^+^ and 4-1BB^+^ T cells (using magnetic beads) (Bacher et al., 2013) or cultured ELISpot assay (Reece et al., 2004) are needed to capture preexisting vaccine-specific memory CD4 T cells directly from human peripheral blood. Another plausible explanation is that our activation proteins, CD40L and 4-1BB, might be unsuitable to detect preexisting memory CD4 T cells but this is unlikely as both proteins have been used successfully in similar studies (Bacher et al., 2014b, 2014a). A different explanation may be that the diversity of preexisting antigen-specific CD4 T clonotypes as determined by TCRβ sequencing is not reflected in the quantitative measurements of the fractional cell counts. A similar disconnect between clonotype diversity underlying vaccine response and cell counts in an ELISPOT setting was observed in Galson et al (Galson et al., 2016). T_REG_ cells represent about 5 – 10% of human CD4 T cell compartment and are identified by the constitutive surface expression of CD25, also known as IL-2 receptor α subunit (IL-2Ra), and the nuclear expression of forkhead family transcription factor 3 (Foxp3), a lineage specification factor of T_REG_ cells (Rudensky, 2011). Regulatory memory T cells play a role in the mitigation of tissue damage induced by effector memory T cells during protective immune responses, resulting in a selective advantage against pathogen-induced immunopathology (Garner-Spitzer et al., 2013; Lanteri et al., 2009; Lin et al., 2018; Lovelace and Maecker, 2018). Several studies have identified CD4 T_REG_ cells with specificity to pathogen-derived peptides in murine models and showed evidence for an induced expansion of T_REG_ cells followed by an emergence and a long-term persistence of T_REG_ cells with a memory phenotype and potent immunosuppressive properties (Lin et al., 2018; Sanchez et al., 2012). Blom et al. reported a significant and transient activation of T_REG_ cells (identified by upregulation of CD38 and Ki67) in humans 10 days after administration of live attenuated yellow fever virus 17D vaccine (Blom et al., 2013). The induction of vaccine-specific T_REG_ cells in our cohort is unexpected and the role it might play in vaccine-induced immunity warrants further investigation.

The association of an expanded 4-1BB^+^ CD45RA^−^ T_REG_ subset with a delayed immune response to hepatitis B vaccine was not described before. Miyara et al. showed that blood contains two distinct subsets of stable and suppressive T_REG_ cells: resting T_REG_, identified as FOXP3^low^CD45RA^+^ CD4 T cells, and activated T_REG_, identified as FOXP3^high^CD45RA^−^ CD4 T cells. They further noted that activated T_REG_ cells constitute a minority subset within cord blood T_REG_ cells and increase gradually with age (Miyara et al., 2009). As activated T_REG_ cells were shown to have an increased expression of proteins indicative of activation, including ICOS and HLA-DR (Booth et al., 2010; Ito et al., 2008; Mason et al., 2015; Miyara et al., 2009; Mohr et al., 2018), it might be the case that an upregulation of 4-1BB is one more feature of this population or a subset thereof. Moreover, T_REG_ cells in mice were shown to modulate T_FH_ formation and GC B cell responses and to diminish antibody production in a CTLA-4 mediated suppression (Wing et al., 2014). Interestingly, CD45RA^−^ T_REG_ cells were shown to be more rich in preformed CTLA-4 stored in intracellular vesicles compared to CD45RA^+^ T_REG_ cells (Miyara et al., 2009).

4-1BB was shown to be constitutively expressed by T_REG_ cells (McHugh et al., 2002) and that 4-1BB^+^ T_REG_ cells are functionally superior to 4-1BB^−^ T_REG_ cells in both contact-dependent and contact-independent immunosuppression (Kachapati et al., 2012). 4-1BB^+^ T_REG_ cells are the major producers of the alternatively-spliced and soluble isoform of 4-1BB among T cells (Kachapati et al., 2012). 4-1BB was shown before to be preferentially expressed on T_REG_ cells compared with other non-regulatory CD4 T cell subsets (McHugh et al., 2002) and that 4-1BB-costimualtion induces the expansion of T_REG_ cells both in vitro and in vivo (Zheng et al., 2004). Moreover, agonistic anti-4-1BB mAbs have been shown to abrogate T cell-dependent antibody responses in vivo (Mittler et al., 1999) and to ameliorate experimental autoimmune encephalomyelitis by skewing the balance against T_H_17 differentiation in favor of T_REG_ differentiation (Kim et al., 2011). It is plausible that the expansion 4-1BB^+^CD45RA^−^ T_REG_ cells in late-converters is involved in the suppression of GC vaccine-specific T_FH_ cells and the ensuing antibody response in our cohort, but this remains speculative and further research is warranted.

It is enticing to speculate that the preexisting memory CD4 T cells result from the complex interplay between cellular immunity and the human microbiome. A role for the microbiota in modulating immunity to viral infection was suggested in 1960’s (Robinson and Pfeiffer, 2014), and since then we gained better understanding of the impact of the various components of the microbiota including bacteria, fungi, protozoa, archaea and viruses on the murine and human immune systems (Winkler and Thackray, 2019).

Viral clearance of hepatitis B virus infection depends on the age of exposure and neonates and young children are less likely to spontaneously clear the virus (Yuen et al., 2018). Han-Hsuan et al. have shown evidence in mice that this age-dependency is mediated by gut microbiota that prepare the liver immunity system to clear HBV, possibly via a TLR4 signaling pathway (Chou et al., 2015). In this study, young mice that have not reached an equilibrium in the gut microbiota, exhibited prolonged HBsAg persistence, impaired anti-HBs antibody production, and limited Hepatitis B core antigen (HBcAg)-specific IFNγ^+^ splenocytes. More recently, Tingxin et al. provided evidence for a critical role of the commensal microbiota in supporting the differentiation of GC B cells, through follicular T helper (T_FH_) cells, to promote the anti-HBV humoral immunity (Wu et al., 2019).

Our study bears some intrinsic limitations. A major drawback is the restricted number of days at which TCRβ repertoire was profiled, as vaccine-specific perturbations within the repertoire may occur at different time points for early, late and non-converters. Additionally, more in-depth characterization and functional studies on 4-1BB^+^CD45RA^−^ T_REG_ cells could have helped shed more light on the role they play in vaccine-induced immunity. Future studies in larger cohorts and with a more comprehensive TCRβ repertoire profiling and CD4 T cells immunophenotyping are required to validate our findings.

In conclusion, our analysis of the memory CD4 T cell repertoire has uncovered a role for preexisting memory CD4 T cells in naïve individuals in mounting an earlier and more vigorous immune response to hepatitis B vaccine and argue for the utility of pre-vaccination TCRβ repertoire in the prediction of vaccine-induced immunity. Moreover, we identify a subset of 4-1BB^+^ memory T_REG_ cells that is expanded in individuals with delayed immune response to the vaccine, which might further explain the heterogeneity of response to hepatitis B vaccine.

## AUTHOR CONTRIBUTIONS

Conception: PM, BO

Design: GE, PM, EB, AS, GM, PVD, PB, KL, VVT, BO

Experiments: GE, PM, EB, NDN, HJ, AS

Data-analysis: GE, PM, NDN, BO

Supervision: HDR, EL, PT, GM, PVD, PB, KL, VVT, BO

First draft: GE, PM, BO

Contributed to the paper: all authors

## Methods

### Human study design and clinical samples

A total of 34 healthy individuals (20-29y: 10, 30-39y: 7, 40-49y: 16, 50+y: 1) without a history of HBV infection or previous hepatitis B vaccination were recruited in this study after obtaining written informed consent. Individuals were vaccinated with a hepatitis B vaccine by intramuscular (m. deltoideus) injection (Engerix-B^®^ containing 20 μg dose of alum-adjuvanted hepatitis B surface antigen, GlaxoSmithKline) on days 0 and 30 (and on day 365). At days 0 (pre-vaccination), 60, 180 and 365, peripheral blood samples were collected on spray-coated lithium heparin tubes, spray-coated K2EDTA (dipotassium ethylenediamine tetra-acetic acid) tubes and serum tubes (Becton Dickinson, NJ, USA).

### Peripheral blood mononuclear cells

Peripheral blood mononuclear cells (PBMC) were isolated by Ficoll-Paque Plus gradient separation (GE Healthcare, Chicago, IL, USA). Cells were stored in 10% dimethyl sulfoxide in fetal bovine serum (Thermo Fisher Scientific, Waltham, MA, USA). After thawing and washing cryopreserved PBMC, cells were cultured in AIM-V medium that contained L-glutamine, streptomycin sulfate at 50 µg/ml, and gentamicin sulfate at 10 µg/ml (Thermo Fisher Scientific, Waltham, MA, USA) and supplemented with 5% human serum (One Lambda, Canoga Park, CA, USA).

### Serology and complete blood count

Serum was separated and stored immediately at – 80°C until time of analysis. Anti-HBs antibody was titrated in serum from day 0, 60, 180 and 365 using Roche Elecsys^®^ Anti-HBs antibody assay on an Elecsys® 2010 analyzer (Roche, Basel, Switzerland). An anti-HBs titer above 10 IU/ml was considered protective (Keating and Noble, 2003).

Serum IgG antibodies to Cytomegalovirus (CMV), Epstein–Barr virus viral-capsid antigen (EBV-VCA), and Herpes Simplex virus (HSV)-1 and 2 were determined using commercially available sandwich ELISA kits in accordance with the manufacturer’s instructions. A complete blood count including leukocyte differential was run on a hematology analyzer (ABX MICROS 60, Horiba, Kyoto, Japan).

### Sorting of memory CD4 T cells

Total CD4 T cells were isolated by positive selection using CD4 magnetic microbeads (Miltenyi Biotech, Bergisch Gladbach, Germany). Memory CD4 T cells were sorted after gating on single viable CD3^+^CD4^+^CD8^−^CD45RO^+^ cells. The following fluorochrome-labeled monoclonal antibodies were used for staining: CD3-PerCP (BW264/56) (Miltenyi Biotech), CD4-APC (RPA-T4) and CD45RO-PE (UCHT1) (both from Becton Dickinson, Franklin Lakes, NJ, USA) and CD8-Pacific Orange (3B5) (from Thermo Fisher Scientific, Waltham, MA, USA). Cells were stained at room temperature for 20 min and sorted with FACSAria II (Becton Dickinson). Sytox blue (Thermo Fisher Scientific) was used to exclude non-viable cells.

### Single peptides, matrix peptide pools and epitope mapping

A set of 15-mers peptides with an 11-amino acid overlap spanning the 226 amino acids along the small S protein of hepatitis B (HB) surface antigen (HBsAg), also designated as small HBs (SHBs) (Shouval, 2003), were synthesized by JPT Peptide Technologies (Berlin, Germany). The set, composed of 54 **single peptides** (See supplementary table 1), was used in a matrix-based strategy to map epitopes against which the immune response is directed (Precopio et al., 2008). The matrix layout enables efficient identification of epitopes within the antigen using a minimal number of cells. For this purpose, a matrix of 15 pools, 7 rows and 8 columns, referred to as **matrix peptide pool**, was designed so that each peptide is in exactly one row-pool and one column-pool, thereby allowing for the identification of positive peptides at the intersection of positive pools. Matrix peptide pools that induced a CD4 T cell response (as determined by CD40L/CD154 assay described below) which meets the threshold criteria for a positive response were considered in the deconvolution process. Top six single peptides were considered for peptide-specific T cell expansion and sorting. A **master peptide pool** is composed of all of the 54 single peptides and was used to identify and sort total vaccine-specific CD4 T cells. Each peptide was used at a final concentration of 2 µg/ml.

### *Ex vivo* T cell stimulation (CD40L/CD154 assay)

Thawed PBMC from each vaccinee were cultured in AIM-V medium that contained L-glutamine, streptomycin sulphate at 50 µg/ml, and gentamicin sulphate at 10 µg/ml. (GIBCO, Grand Island, NY) and supplemented with 5% human serum (One Lambda, Canoga Park, CA, USA). Cells were stimulated for 6 hours with 2 μg/ml of each of the 15 matrix peptide pools in the presence of 1 µg/ml anti-CD40 antibody (HB14) (purchased from Miltenyi Biotec, Bergisch Gladbach, Germany) and 1 μg/ml anti-CD28 antibody (CD28.2) (purchased from BD Biosciences, Franklin Lakes, NJ, USA).

Cells were stained using the following fluorochrome-labelled monoclonal antibodies: CD3-PerCP (BW264/56), CD4-APC (REA623), CD8-VioGreen (REA734) and CD40L-PE (5C8) (purchased from Miltenyi Biotec, Bergisch Gladbach, Germany). Viability dye Sytox blue from Invitrogen (Thermo Fisher Scientific, Waltham, MA, USA) was used to exclude non-viable cells.

Data was acquired on FACSAria II using Diva Software, both from BD Biosciences (Franklin Lakes, NJ, USA), and analyzed on FlowJo software version 10.5.3 (Tree Star, Inc., Ashland, OR, USA). Fluorescence-minus-one controls were performed in pilot studies. Gates for CD40L^+^CD4 T cells were set using cells left unstimulated.

### *In vitro* T cell expansion and cell sorting

Thawed PBMC were labelled with carboxyfluorescein succinimidyl ester (CFSE) (Invitrogen, Carlsbad, CA, USA) and cultured in AIM-V medium that contained L-glutamine, streptomycin sulphate at 50 µg/ml, and gentamicin sulphate at 10 µg/ml. (GIBCO, Grand Island, NY) and supplemented with 5% human serum (One Lambda, Canoga Park, CA, USA). Cells were stimulated for 7 days with 2 µg/ml of selected single peptides in addition to the master peptides pool. Cells were stained using the following fluorochrome-labelled monoclonal antibodies: CD3-PerCP (BW264/56), CD4-APC (REA623) and CD8-VioGreen (REA734) (purchased from Miltenyi Biotec, Bergisch Gladbach, Germany). Viability dye Sytox blue from Invitrogen (Thermo Fisher Scientific, Waltham, MA, USA) was used to exclude non-viable cells. Single viable CFSE^low^ CD3^+^ CD8^−^ CD4^+^ T cells were sorted into 96-well PCR plates containing DNA/RNA Shield (Zymo Research, Irvine, CA, USA) using FACSAria II and Diva Software (BD Biosciences, Franklin Lakes, NJ, USA). For each of the selected single peptides, 500 cells were sorted in two technical replicates. For the master peptide pool, 1000 cells were sorted in two technical replicates. Plates were immediately centrifuged and kept at – 20°C before TCR cDNA library preparation and sequencing.

### TCRβ cDNA Library Preparation and Sequencing of memory CD4 T cells

DNA was extracted from sorted memory CD4 T cells using Quick-DNA Microprep kit (Zymo Research, Irvine, CA, USA). ImmunoSEQ hsTCRB sequencing kit (Adaptive Biotechnologies, Seattle, WA, USA) was used to profile TCRβrepertoire following the manufacturer’s protocol.

After quality control using Fragment Analyzer (Agilent, Santa Clara, CA, USA), libraries were pooled with equal volumes. The concentration of the final pool was measured with the Qubit™ dsDNA HS Assay kit (Thermo Fisher Scientific, Waltham, MA, USA). The final pool was processed to be sequenced on the Miseq and NextSeq platforms (Illumina, San Diego, CA, USA). Memory CD4 T cells of one of the vaccinees (H42, a non-converter) was not sequenced due to a capacity issue.

### TCR cDNA Library Preparation and Sequencing of CFSE^low^ CD4 T cells

RNA was extracted from each of the two technical replicates of sorted CFSE^low^ CD4 T cells using Quick-RNA Microprep kit (Zymo Research, Irvine, CA, USA). Without measuring the resulted RNA concentration, an RNA-based library preparation was used. The QIAseq Immune Repertoire RNA Library kit (Qiagen, Venlo, Netherlands) amplifies TCR alpha, beta, gamma and delta chains. After quality control using Fragment Analyzer (Agilent, Santa Clara, CA, USA), concentration was measured with the Qubit™ dsDNA HS Assay kit (Thermo Fisher Scientific, Waltham, MA, USA) and pools were equimolarly pooled and prepared for sequencing on the Nextseq platform (Illumina, San Diego, CA, USA).

### TCRβ Sequence Analysis

TCRβ clonotypes were identified as previously described (de Neuter et al., 2018) where a unique TCRβ clonotype is defined as a unique combination of a V gene, CDR3 amino acid sequence, and J gene. All memory CD4 T cell DNA-based TCRβ sequencing reads were annotated using the immunoSEQ analyzer (v2) from Adaptive Biotechnologies. All small bulk RNA-based TCR sequencing reads were annotated using the MiXCR tool (v3.0.7) from the FASTQ files. As all RNA-based TCR sequencing experiments featured two technical replicates, only those TCR sequences that occurred in both replicates were retained and their counts were summed. Tracking of vaccine-specific TCRβ clonotypes is based on exact TCRβ CDR3 amino acid matches to remove any bias introduced by the different VDJ annotation pipelines. Non-HBsAg TCR annotations were done with the TCRex web tool (Gielis et al., 2019) on the 24^th^ of July, 2019 using version 0.3.0. Inference of similar epitope binding between two TCR sequences is defined according to the Hamming distance (*d*) calculated on the CDR3 amino acid sequence with a cutoff *c*, as supported by Meysman et al. (Meysman et al., 2018). All scripts used in this analysis are available via github (https://github.com/pmeysman/HepBTCR).

### Predictive HBs-response model

From the single peptide data generated in the matrix peptide pool experiments, we aimed to create a predictive model to enumerate the HBs response from full TCRβ repertoire data. This approach allows for predictions that are epitope-specific rather than simply vaccine-specific. This model was applied in a leave-one-out cross validation so that vaccine-specific TCRβ sequences from a vaccinee are not used to make predictions for the same vaccinee. While the predictive model is derived from epitope-specific data, it cannot be guaranteed that some of the expanded CD4 T cells detected in the in vitro assay are not due to bystander activation. Vaccine-specific TCRβ sequences of vaccinees who did not respond to the vaccine at day 60 (late-converters and non-converters) are expected to be more enriched in cells triggered to expand due to bystander activation. Indeed, running the set of vaccine-specific TCRβ sequences through the TCRex webtool (Gielis et al., 2019) reveals that several TCRs are predicted to be highly similar to those reactive to the CMV NLVPMVATV epitope (enrichment P value <0.001 when compared to the TCRex background repertoire) and the Mart-1 variant ELAGIGILTV epitope (P value <0.001), which supports the notion that some of these TCRβ sequences might not be specific to HBsAg. This set of vaccine-specific TCRβ sequences can thus be used to make predictions about possible TCRβ sequences due to bystander activation of CD4 T cells, i.e. common TCRβ sequences that might be present as false positives. The final output of the model is thus a ratio, **R***_hbs_*, for any repertoire rep*_i_* describing a set of TCRβ sequences *t_repi_*:

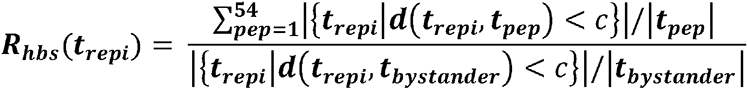

with *t_pep_* as the set of TCRβ sequences occurring in both biological replicates for a single sample and a single peptide (*pep*) from the HBsAg matrix peptide pool experiment, and *t_bystander_* as the set of TCRβ sequences occurring in both biological replicates of the master peptide pool in any of the non-responding samples. Thus the ratio signifies the number of TCR clonotypes predicted to be reactive against one of the HBsAg peptides, normalized by a count of putative false positive predictions from bystander T-cells.

### *Ex vivo* T cell phenotyping of vaccine-specific T cells

Thawed PBMC from each vaccinee were cultured in AIM-V medium that contained L-glutamine, streptomycin sulphate at 50 µg/ml, and gentamicin sulphate at 10 µg/ml. (GIBCO, Grand Island, NY) and supplemented with 5% human serum (One Lambda, Canoga Park, CA, USA). Cells were stimulated for 6 hours with 2 μg/ml of a master peptide pool representing the full length of the small surface envelope protein of hepatitis B, in the presence of 1 µg/ml anti-CD40 antibody (HB14) (purchased from Miltenyi Biotec, Bergisch Gladbach, Germany) and 1 μg/ml anti-CD28 antibody (CD28.2) (purchased from BD Biosciences, Franklin Lakes, NJ, USA). Cells were stained using the following fluorochrome-labelled monoclonal antibodies:

CD3-BV510 (SK7), CD4-PerCP/Cy5.5 (RPA-T4), CD8-APC/Cy7 (SK1), CD45RA-AF488 (HI100), CD25-BV421 (M-A251), CD127-BV785 (A019D5) and CD137-PE (4-1BB) (purchased from BioLegend, San Diego, CA, USA), CXCR5 (CD185)-PE-eFluor 610 (MU5UBEE) (from eBioscience, Thermo Fisher Scientific, Waltham, MA, USA) and CD40L-APC (5C8) (purchased from Miltenyi Biotec, Bergisch Gladbach, Germany). Fixable viability dye Zombie NIR™ from BioLegend (San Diego, CA, USA) was used to exclude non-viable cells. Data was acquired on FACSAria II using Diva Software, both from BD Biosciences (Franklin Lakes, NJ, USA), and analyzed on FlowJo software version 10.5.3 (Tree Star, Inc., Ashland, OR, USA) using gating strategy shown in **Fig. S1a**. Fluorescence-minus-one controls were performed in pilot studies. Gates for CD40L^+^ and 4-1BB^+^ CD4 T cells (**Fig. S1b**) were set using cells left unstimulated (negative control contained DMSO at the same concentration used to solve peptide pools). In order to account for background expression of CD40L and 4-1BB on CD4 T cells, responses in cells left unstimulated were subtracted from the responses to peptides, and when peptides-specific CD40L^+^ or 4-1BB^+^ CD4 T cells were not significantly higher than those detected for cells left unstimulated (using one-sided Fisher’s exact test), values were mutated to zero.

### Statistics and data visualization

The two-sided Fisher’s exact test was used to evaluate the significance of relationship between early/late-converters and CMV, EBV or HSV seropositivity. For the visualization of marker expression, TCRβ counts and cell frequencies between time points or groups of vaccinees, ggplot2 (V3.3.2) and ggpubr (V0.2.5) packages in R were used. The Wilcoxon signed-rank test was used to compare two or more groups, with unpaired and paired analysis as necessary. The nonparametric Spearman’s rank-order correlation was used to test for correlation. We used the following convention for symbols indicating statistical significance; ns P > 0.05, * P ≤ 0.05, ** P ≤ 0.01, *** P ≤ 0.001, **** P ≤ 0.0001.

### Study approval

Protocols involving the use of human tissues were approved by the Ethics Committee of Antwerp University Hospital and University of Antwerp (Antwerp, Belgium), and all of the experiments were performed in accordance with the protocols.

### Data availability

The sequencing data that support the findings of this study have been deposited on Zenodo (https://doi.org/10.5281/zenodo.3989144). Flow Cytometry Standard (FCS) data files with associated FlowJo workspaces are deposited at flowrepository.org (Spidlen et al., 2012) under the following experiment names: epitope mapping: https://flowrepository.org/id/FR-FCM-Z2TN; *in vitro* T cell expansion: https://flowrepository.org/id/FR-FCM-Z2TM; *ex vivo* CD4 T cell assay: https://flowrepository.org/id/FR-FCM-Z2TL.

---

**Figure S1.**
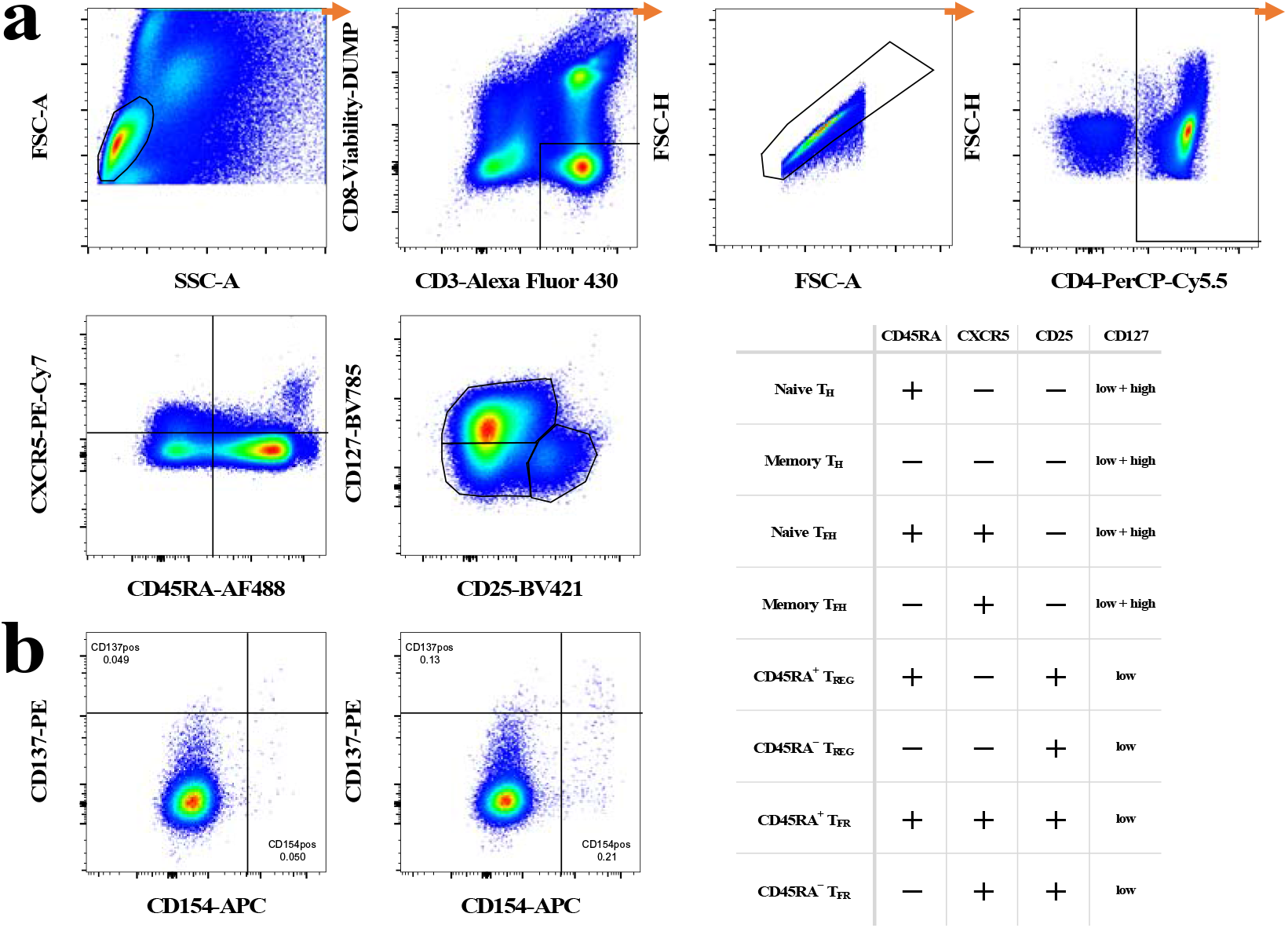
Gating strategy of ex vivo T cell phenotyping of vaccine-specific T cells. **a** Gating strategy started by a lymphocyte gate, followed by gating on viable CD3^+^CD8^−^ T cells. Doublets were excluded using doublet discrimination (area against the height of forward scatter pulse) before gating on CD4^+^ T cells. Next, CD45RA, CXCR5, CD25 and CD127 were used to identify main subsets of CD4 T cells using Boolean gates as specified in the accompanying table. **b** Shown an example of gating for CD154 (CD40L) and CD137 (4-1BB) for cells left unstimulated (left) and cells stimulated with a master peptide pool (right) for an early-converter vaccinee at day 60.

**Figure S2.**
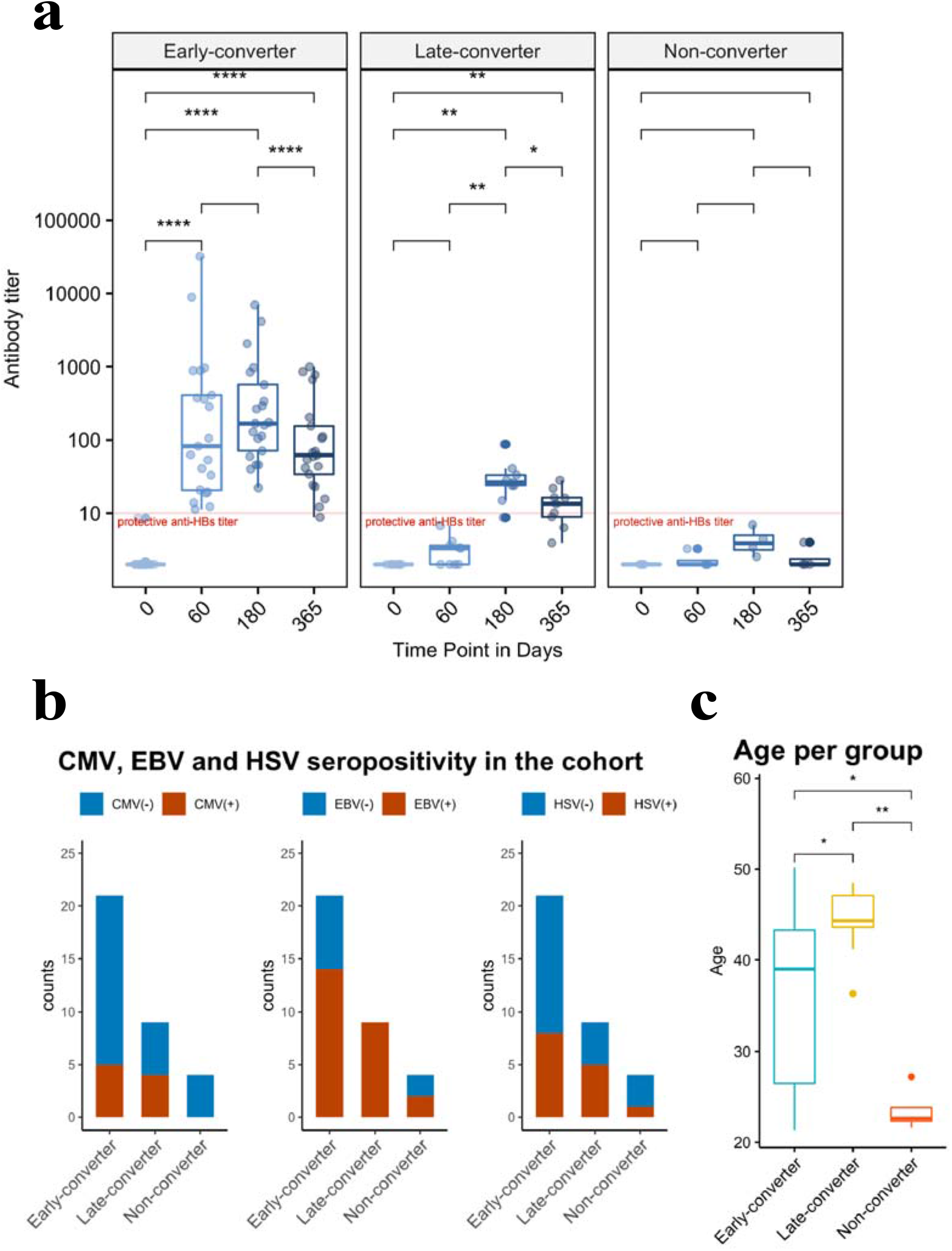
Serological memory to hepatitis B vaccine and vaccinee groups within the cohort. a Anti-Hepatitis B surface (anti-HBs) titer of vaccinees over four times points, facetted by groups of early, late and non-converters. An anti-HBs titer above 10 IU/ml was considered protective. Early-converters seroconverted at day 60, late-converters seroconverted at day 180 or day 365 and non–converters did not have an anti-HBs titer higher than 10 IU/ml at any of the time points. b CMV, EBV and HSV seropositivity in the three groups of the cohort as determined by serum IgG antibodies to CMV, EBV-VCA, and HSV-1 and 2 using sandwich ELISA. c Age of vaccinees per group. Wilcoxon signed-rank with unpaired and paired analysis as necessary; statistical significance was indicated with ns P > 0.05, * P ≤ 0.05, ** P ≤ 0.01, *** P ≤ 0.001, **** P ≤ 0.0001

**Figure S3.**
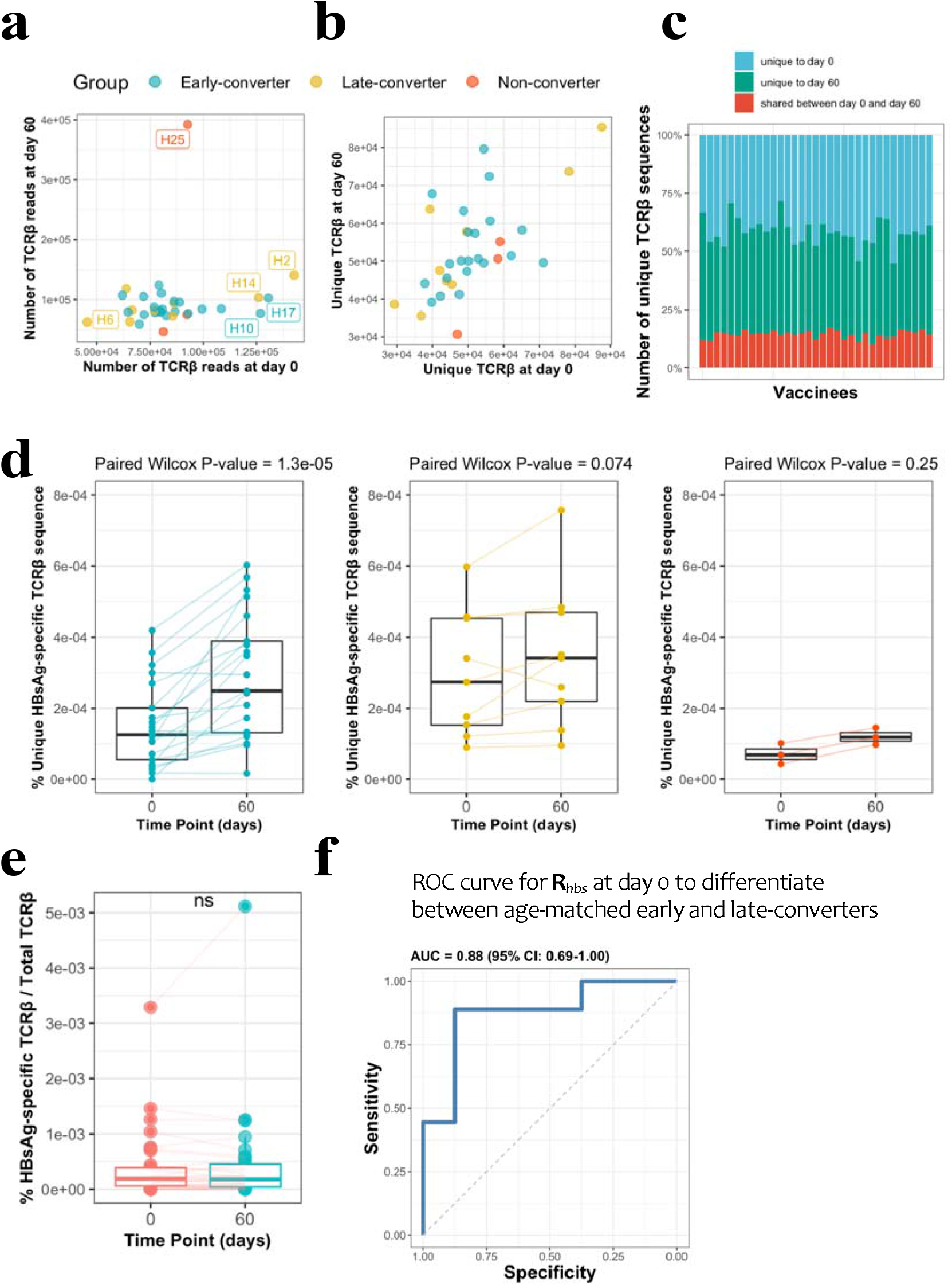
CD4 T cell memory TCRβ repertoire and vaccine-specific TCRβ clonotypes. a Scatter plot of the DNA-based TCRβ reads for each vaccinee at each time point. b Scatter plot of number of unique TCRβ amino acid sequences for each vaccinee at each time point, where the shape denotes the response as based on antibody titer. c Overview of unique TCRβ amino acid sequences in the memory CD4 T cell repertoire of each vaccinee. The bottom blue bar denotes those TCRβ sequences that were found at both time points. The green and red bars denote the number of unique TCRβ sequences at each time point. The total bar height thus represents the total number of unique memory CD4 T cell clonotypes sequences for a specific vaccinee. d Frequency of unique HBsAg-specific TCRβ sequences out of total sequenced TCRβ sequences between two time points for all vaccinees colored and faceted by group. e Change in frequency of those HBsAg-specific CD4 T cells present at both time points. The (ns) mark denotes a non-significant paired Wilcoxon signed-rank test (p-value = 0.7577). f Receiver operating characteristic (ROC) curve using R*_hbs_* to differentiate between age-matched early-converters and late-converters in a leave-one-out cross validation at day 0. Age-matching was accomplished retaining only samples in the age range 40-55. A Wilcoxon test was used to confirm that there was no difference in age distributions between early and late converters (P value = 0.60, mean EC = 44.5y, mean LC 45.1y). Diagonal line denotes a random classifier. Reported is the area under the curve (AUC) and its 95% confidence interval.

**Figure S4.**
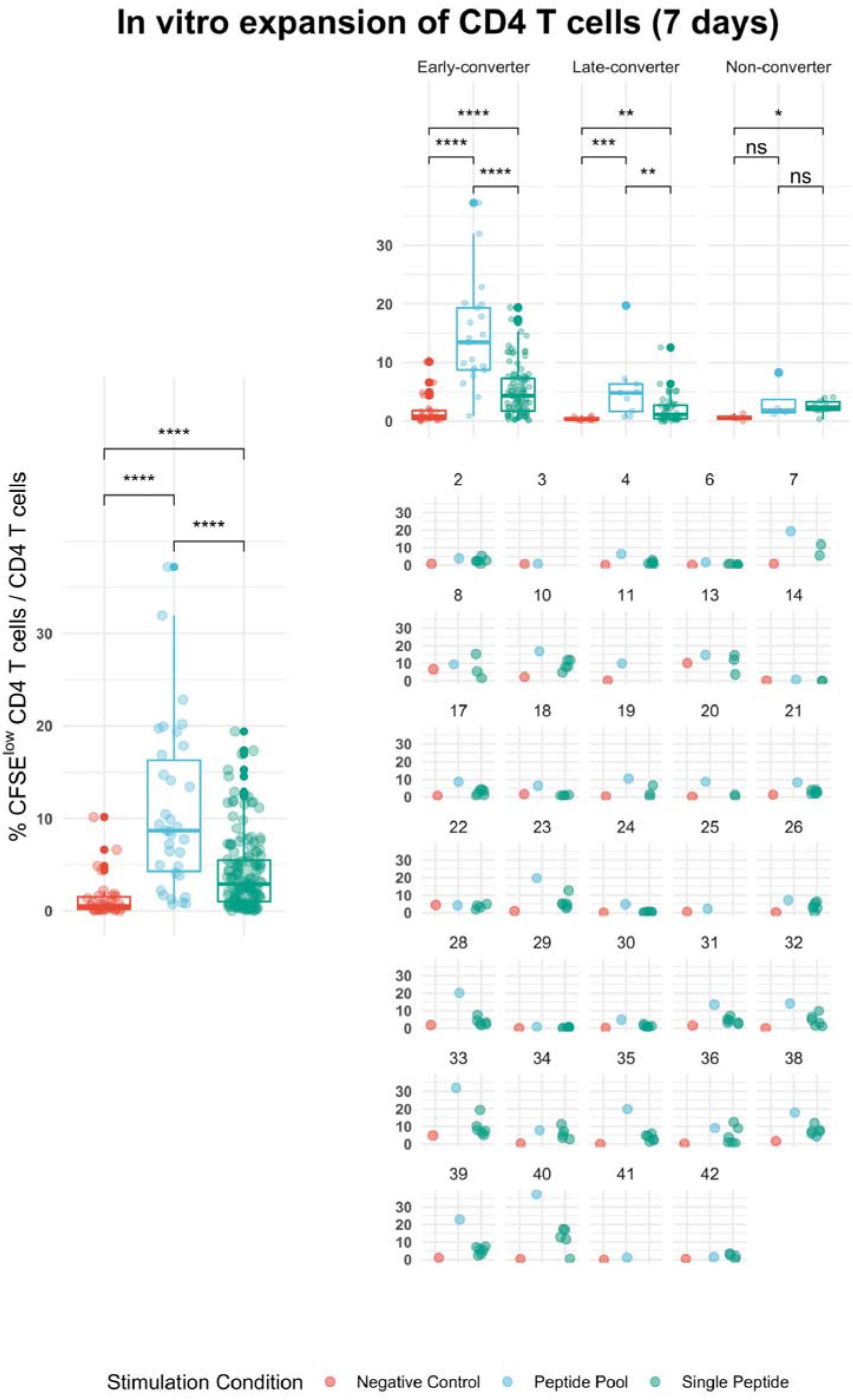
Overview of the results of in vitro expansion experiments. Shown is the frequency of CFSE^low^ CD4 T cells out of total CD4 T cells for all vaccinees, vaccinees per group and for each vaccinee. Peripheral blood mononuclear cells from day 60 were labeled with carboxyfluorescein succinimidyl ester (CFSE) and stimulated with a pool of peptides spanning hepatitis B (HB) surface antigen (HBsAg) (**Peptide Pool**) and single peptides selected based on epitope mapping of the entire antigen (**Single Peptide**). After day 7 of in vitro expansion, cells were stained with antibodies to surface markers (CD3, CD4 and CD8) that enable gating on viable CD4 T cells. CFSE intensity was used to identify and sort CFSE^low^ cells for TCR repertoire analysis of antigen-specific CD4 T cells.

**Figure S5.**
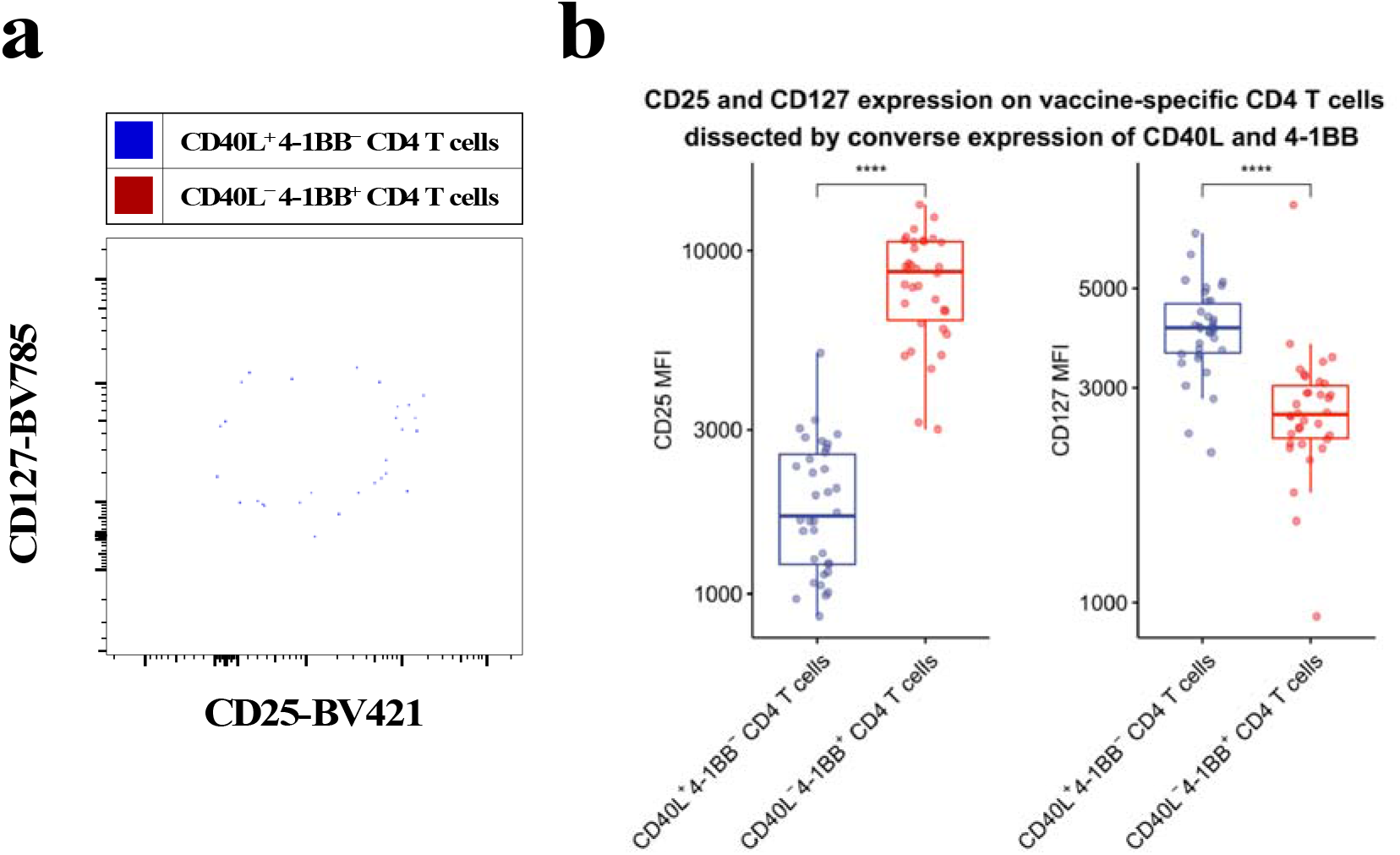
CD40L^+^4-1BB^−^ and CD40L^−^4-1BB^+^ CD4 T cells have a T_CON_ and T_REG_ phenotype, respectively. PBMCs from vaccinees were stimulated with 2 μ ml of a pool of peptides of HBsAg and assessed for converse expression of 4-1BB and CD40L by flow cytometry. a CD40L^+^4-1BB^−^ and CD40L^−^4-1BB^+^ CD4 T cells from day 60 were gated on and then overlaid in a contour plots of CD25 versus CD127 to assess T_COV_ and T_REG_ phenotype. b summary plot of median fluorescence intensity (MFI) of CD25 and CD127 for all vaccinees. Wilcoxon signed-rank with paired analysis; statistical significance was indicated with **** P ≤ 0.0001

**Figure S6.**
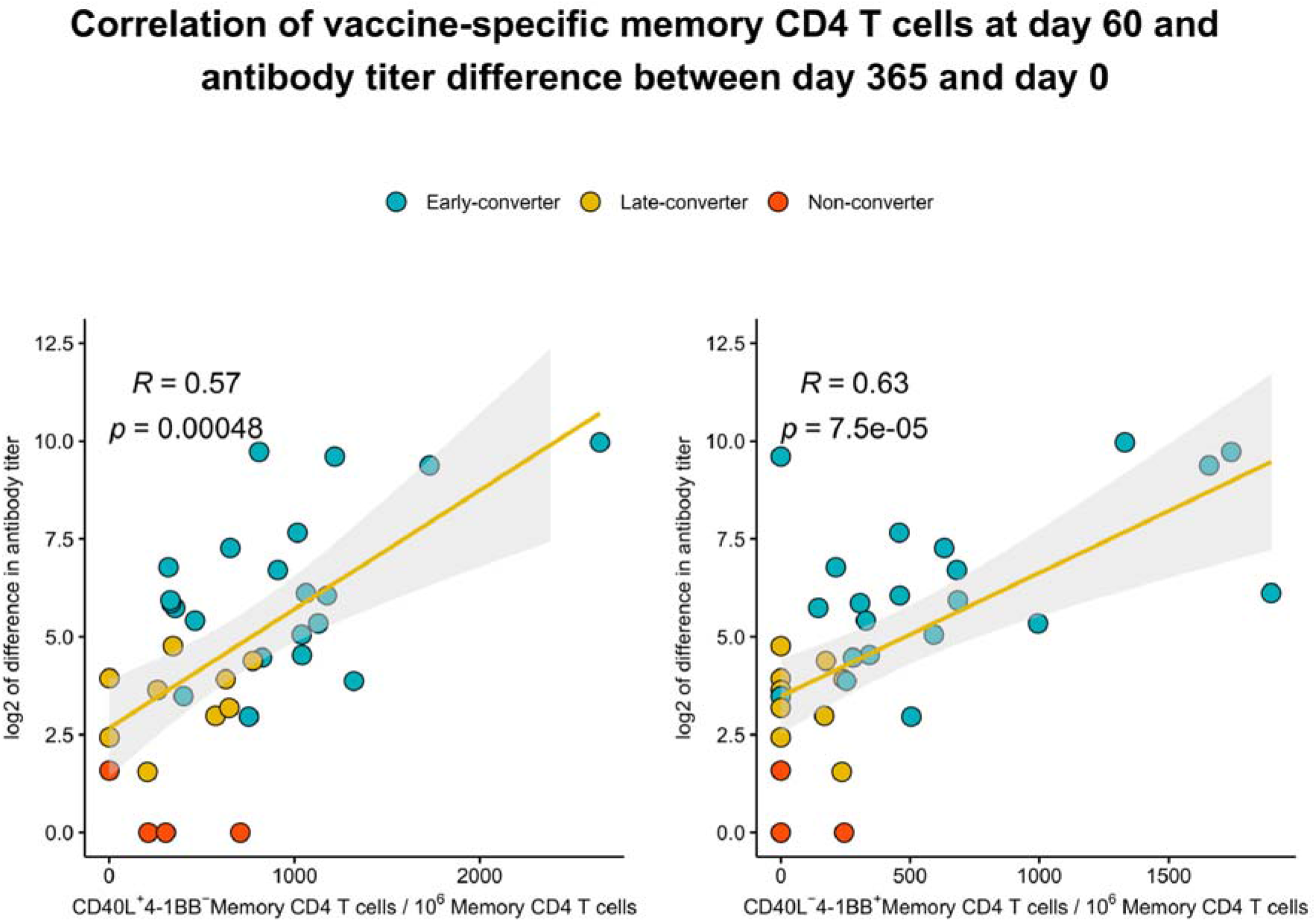
Relationship between serological memory and memory CD4 T cell response to the vaccine. Correlation between the difference in antibody titer between day 365 and day 0 and vaccine-specific CD40L^+^4-1BB^−^ and CD40L^−^4-1BB^+^ memory CD4 T cell at day 60 colored by vaccinee group and labeled with vaccinee ID. *rs*, Spearman correlation coefficient, −1 ≤ *r*s ≤ 1; *rs* and *p* value by Spearman’s correlation test

**Figure S7.**
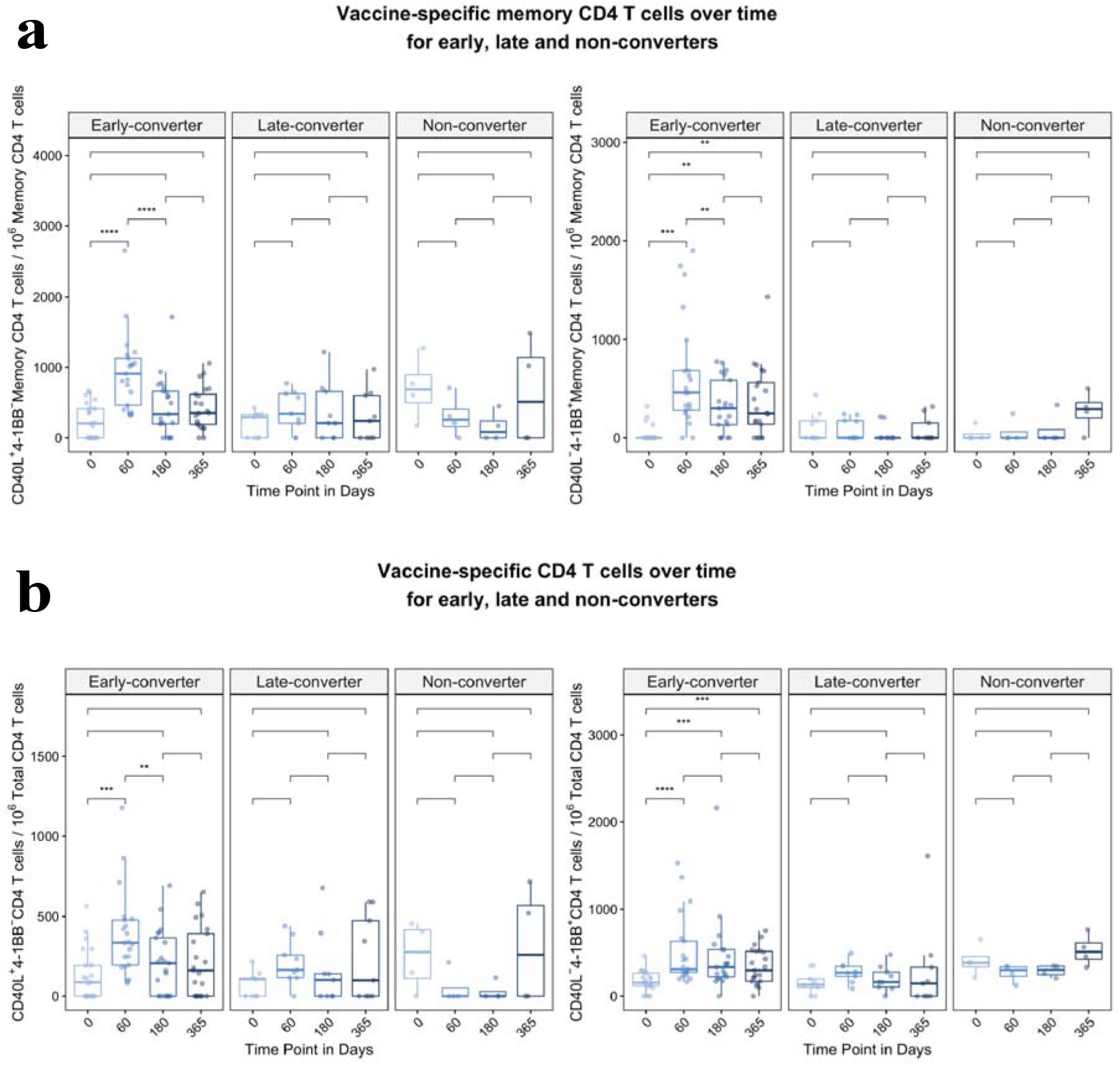
Hepatitis B vaccine induces a vaccine-specific CD4 T cell response in early-converter vaccinees. PBMCs from vaccinees were stimulated with 2 μg/ml of a pool of peptides of HBsAg and assessed for converse expression of 4-1BB and CD40L by flow cytometry on days 0, 60, 180, and 365. **a** Aggregate analysis from early, late and non-converter vaccinees showing a significant induction of vaccine-specific CD40L^+^4-1BB^−^ and CD40L^−^4-1BB^+^ memory CD4 T cell in early-converters and lack thereof in late and non-converters. Shown are numbers of vaccine-specific memory CD4 T cells out of 10^6^ memory CD4 T cells after subtraction of responses in negative control (see Methods for details). **b** Aggregate analysis from early, late and non-converter vaccinees showing a significant induction of vaccine-specific CD40L^+^4-1BB^−^ and CD40L^−^4-1BB^+^ CD4 T cell in early-converters and lack thereof in late and non-converters. Shown are numbers of vaccine-specific CD4 T cells out of 10^6^ CD4 T cells after subtraction of responses in negative control (see Methods for details).

**Figure S8.**
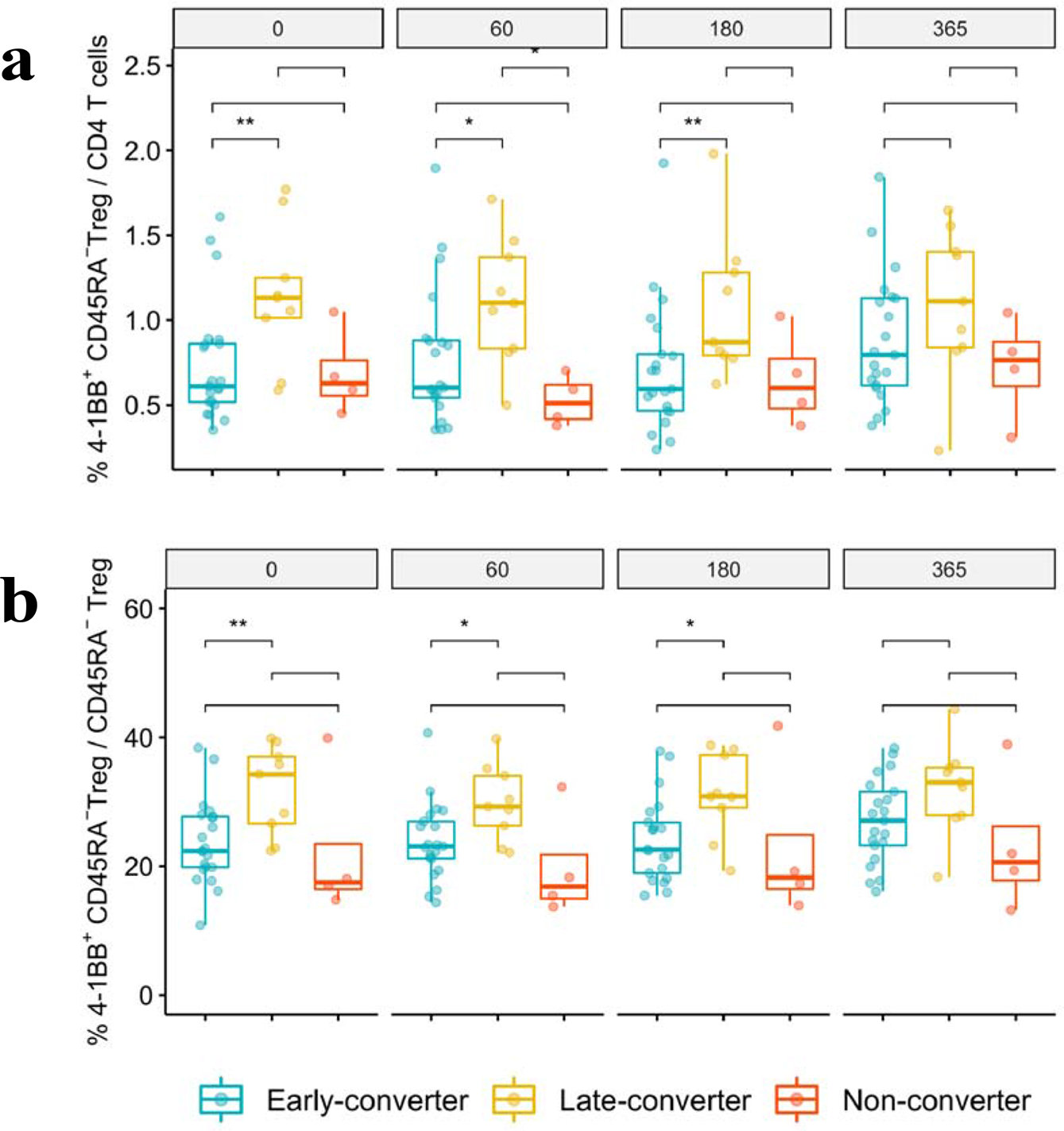
An expanded 4-1BB^+^CD45RA^−^ T_REG_ cells within T_REG_ compartment is a prominent feature in late-converters prior to vaccination. Aggregate analysis of the frequency of 4-1BB^+^CD45RA^−^ T_REG_ within **a** total CD4 T cells and **b** CD45RA^−^ T_REG_ CD4 T cells in early, late and non-converter vaccinees at days 0, 60, 180 and 365. Wilcoxon signed-rank with unpaired and paired analysis as necessary; statistical significance was indicated with ns P > 0.05, * P ≤ 0.05, ** P ≤ 0.01, *** P ≤ 0.001, **** P ≤ 0.0001

## Supplemental tables

**Supplementary Table 1.**
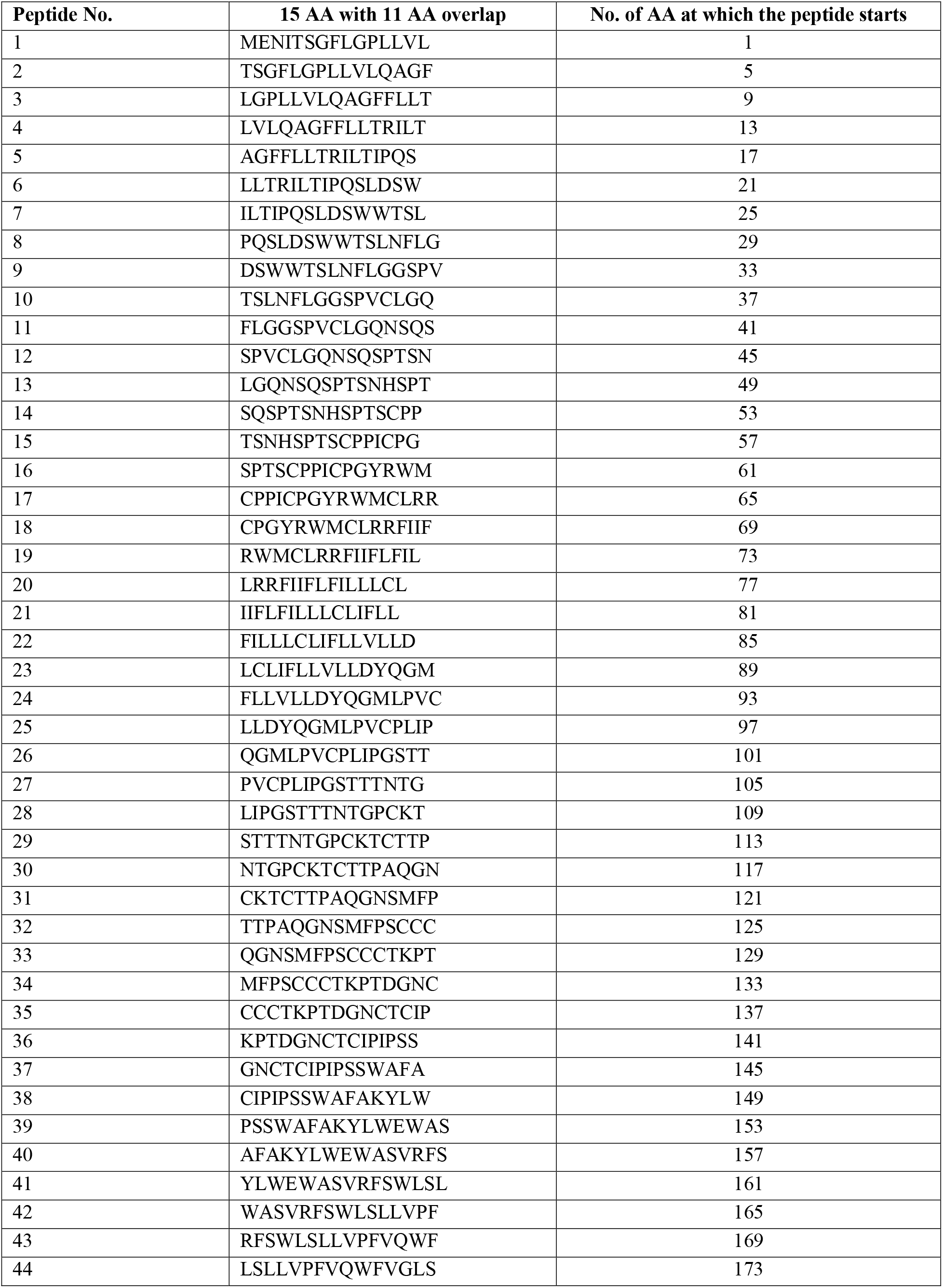
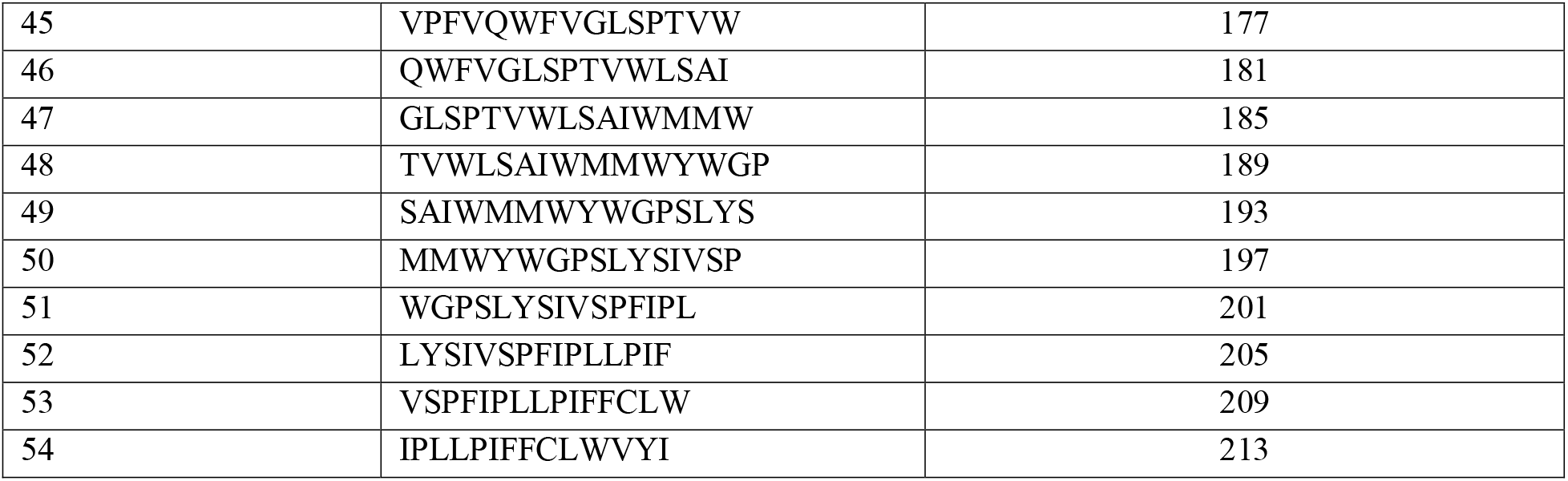
List of 54 single peptides, each 15 AA long with an 11-amino acid overlap spanning the 226 amino acids along the small S protein of hepatitis B (HB) surface antigen (HBsAg)

**Supplementary Table 2.** Overview of the single peptides tested for each vaccinee in the CFSE assay

| Vaccinee | Gender | Age | Status | peptides_number | single peptides |
| --- | --- | --- | --- | --- | --- |
| 2 | F | 43.6 | Late-converter | 6 | 51, 35, 50, 34, 54, 38 |
| 3 | F | 48.5 | Late-converter | 6 | 53, 5, 37, 49, 1, 33 |
| 4 | M | 47.1 | Late-converter | 6 | 21, 20, 5, 53, 4, 52 |
| 6 | F | 44.3 | Late-converter | 6 | 23, 39, 7, 21, 37, 5 |
| 7 | M | 38.3 | Early-converter | 2 | 17, 33 |
| 8 | M | 43.4 | Early-converter | 3 | 47, 41, 42 |
| 10 | F | 33.2 | Early-converter | 5 | 49, 1, 33, 41, 9 |
| 11 | F | 39 | Early-converter | 6 | 53, 50, 5, 2, 8, 21 |
| 13 | F | 45.9 | Early-converter | 3 | 24, 8, 16 |
| 14 | M | 36.3 | Late-converter | 2 | 45, 47 |
| 17 | F | 26.5 | Early-converter | 6 | 10, 42, 9, 41, 14, 46 |
| 18 | M | 43.3 | Early-converter | 4 | 6, 38, 5, 37 |
| 19 | M | 41.3 | Early-converter | 3 | 40, 16, 8 |
| 20 | F | 48.7 | Early-converter | 2 | 6, 1 |
| 21 | F | 27.2 | Non-converter | 6 | 2, 1, 3, 42, 41, 43 |
| 22 | F | 22.3 | Early-converter | 4 | 10, 13, 12, 16 |
| 23 | F | 43.9 | Late-converter | 6 | 51, 54, 48, 43, 47, 46 |
| 24 | F | 47.7 | Late-converter | 6 | 22, 24, 23, 20, 18, 46 |
| 25 | F | 22.6 | Non-converter | 6 | 29, 30, 28, 21, 22, 20 |
| 26 | F | 41.2 | Late-converter | 6 | 23, 47, 17, 41, 7, 49 |
| 28 | F | 41.7 | Early-converter | 6 | 54, 38, 52, 36, 40, 39 |
| 29 | M | 39.5 | Early-converter | 6 | 7, 47, 6, 1, 23, 46 |
| 30 | F | 45.1 | Late-converter | 6 | 22, 46, 54, 19, 20, 43 |
| 31 | M | 32.4 | Early-converter | 6 | 54, 38, 49, 33, 53, 37 |
| 32 | F | 31.4 | Early-converter | 6 | 7, 4, 1, 6, 31, 28 |
| 33 | F | 41 | Early-converter | 6 | 1, 41, 7, 5, 47, 45 |
| 34 | M | 22.5 | Early-converter | 5 | 7, 4, 5, 2, 6 |
| 35 | F | 50.2 | Early-converter | 6 | 46, 45, 47, 22, 21, 23 |
| 36 | F | 23.2 | Early-converter | 6 | 54, 47, 46, 39, 48, 38 |
| 38 | F | 23.2 | Early-converter | 6 | 37, 34, 33, 35, 5, 2 |
| 39 | M | 45.8 | Early-converter | 6 | 51, 50, 53, 49, 35, 34 |
| 40 | F | 21.3 | Early-converter | 5 | 33, 38, 37, 39, 40 |
| 41 | M | 21.6 | Non-converter | 6 | 23, 22, 7, 6, 17, 1 |
| 42 | M | 22.7 | Non-converter | 4 | 1, 33, 25, 41 |

